# Microtubule curling as an efficient readout to uncover fundamental concepts of axonal cell biology and neurodegeneration

**DOI:** 10.1101/2025.06.27.662005

**Authors:** André Voelzmann, Milli Owens, Robin Beaven, William Cairns, Abigail Elliot, Sheng-Hui Feng, Judith Fuelle, Catarina Goncalves-Pimentel, Ines Hahn, Ella Jones, Kodie Norris, Yu-Ting Liew, Lydia Lorenzo-Cisneros, Lucy O’Donoghue, Liliana M. Pinho-Correia, Ananya Pupala, Yue Qu, Tarunima Sharma, Natalia Sánchez-Soriano, Andreas Prokop

## Abstract

Nerve fibres (aka axons) are the slender, up-to-meter-long processes of nerve cells that wire nervous systems. These delicate structures must survive for an organism’s lifetime, making them prime lesion sites in neurodegeneration. Their long-term maintenance requires the homeostasis of complex local cell biology upheld by motor protein-driven transport along microtubule bundles that run uninterrupted along axons. To gain an understanding of axonal homeostasis, we report and discuss here the functional loss of 105 genes from a wide range of cell biological processes using a standardised *Drosophila* primary neuron system. ∼40% of these gene deficiencies caused microtubule bundle disturbances referred to as ‘microtubule-curling’, which we use as an indicator of axonal atrophy. Our live imaging showed that microtubule-curling initiates in areas where axons widen, such as areas close to the soma, branch points or growth cones. In wild-type neurons, any initiated curling was contained, but it persisted as growing footprints in mutant curl-promoting conditions. Closer analyses of 20 curl-promoting conditions suggested a general classification into two groups: mutations affecting axonal physiology cause ROS-mediated microtubule-curling, whereas mutations causing motor hyperactivation or affecting microtubule-regulating proteins cause structurally induced ROS-independent curling. As will be discussed, our data provide consistent support for the previously proposed ‘dependency cycle of local axon homeostasis’ model which can explain the long-standing conundrum that genes from a wide range of cell biological processes often have mutational links to the same class of inherited neurodegenerative disease.

## Introduction

Neurodegenerative disorders are an increasing socioeconomic burden to modern ageing societies causing enormous suffering to affected individuals (Steinmetz et al., 2024). However, due to the enormous complexity of neuronal cell biology, the underlying pathological processes remain little understood posing an obstacle to finding cures. A potential way to narrow down the task is to focus on nerve fibres, aka axons. Axons are the up-to-meter-long cable-like protrusions of neurons that mediate the crucial information flow of nervous systems. Despite being exposed to constant intra- and extracellular mechanical challenges, these delicate structures can usually not be replaced and must survive for an organism’s lifetime (i.e. up to a century in humans); consequently, ∼40% of axons are lost during normal ageing in mammals (Adalbert and Coleman, 2012; Calkins, 2013; Marner et al., 2003), and they are prime lesion sites in many neurodegenerative diseases (Coleman and Höke, 2020; Prokop, 2021a; Smith et al., 2023). A key question therefore is how axons can survive long-term.

Since distal portions of axons can be meters away from their associated cell bodies, their long-term maintenance requires the local presence and homeostasis of essential cell biological components and functions including organelles, endomembrane systems, cytoskeleton, metabolism, proteostasis and signalling mechanisms, all jointly arranged into a network of local homeostasis (Smith et al., 2023). Maintaining this homeostasis requires the transport and movement of organelles and materials from and to the cell body or, locally, to sites of acute need; this transport is mediated by molecular motors which move along microtubules (MTs). For this, axonal MTs are arranged into bundles that serve as essential transport highways by running uninterrupted from the neuronal cell body to the axon tip (Guedes-Dias and Holzbaur, 2019; Prokop, 2020). In this way, MT bundles throw a lifeline to axons which, if disrupted, can also become their Achilles heels.

Using *Drosophila* neurons as a standardised cellular model to study the role and regulation of MTs in the context of axonal cell biology, we generated data that led us to propose the ‘dependency cycle of local axon homeostasis’ as a model that may help to conceptualise processes of axonal degeneration (Fig.1A;)(Liew et al., 2026; Prokop, 2021a; Smith et al., 2023). The dependency cycle proposes that axonal transport is beneficial for axon maintenance (‘1’ and ‘2’ in Fig.1A). But transport also imposes mechanical forces (‘3’) that can lead to MT bundle disorganisation (referred to as ‘MT-curling’; Fig.1B-G) potentially triggered through buckling (Fig.2;)(Bicek et al., 2009), physical damage that weakens the MT lattice (Dumont et al., 2015), or deflecting the extension of polymerising MTs out of their parallel bundle arrangements (‘2’ in Fig.2). Like potholes in our streets are kept in check by highway maintenance, MT-curling is counter-balanced by MT-binding proteins that protect, repair or replace MTs and their organisation (‘4’; examples in Fig.2), thus establishing ‘structural homeostasis’ (left green box in Fig.1A). In this scenario, loss of axonal transport may bring structural relief, but it also impairs cargo availability and disturbs ‘physiological homeostasis’ (right green box) which, in-turn, triggers ROS dyshomeostasis as a powerful inducer of MT-curling (‘ROS’ in Fig.1A;)(Liew et al., 2026; Shields et al., 2025). Taken together, the model proposes that structural homeostasis is required upstream of proper axonal transport, but it is also downstream of and dependent on functioning axonal transport and physiological homeostasis (‘5’) – thus establishing a circular relationship. We predict therefore that the homeostasis of this circular relationship can be broken by the dysfunction of very different aspects of axonal cell biology, which should then be reflected in MT-curling as a common pathology indicator. If correct, our model might provide an explanations for the conundrum that mutations in a wide range of genes often link to the same class of neurodegenerative disorder (Prokop, 2021a; Shields et al., 2024; Smith et al., 2023).

**Fig. 1.**
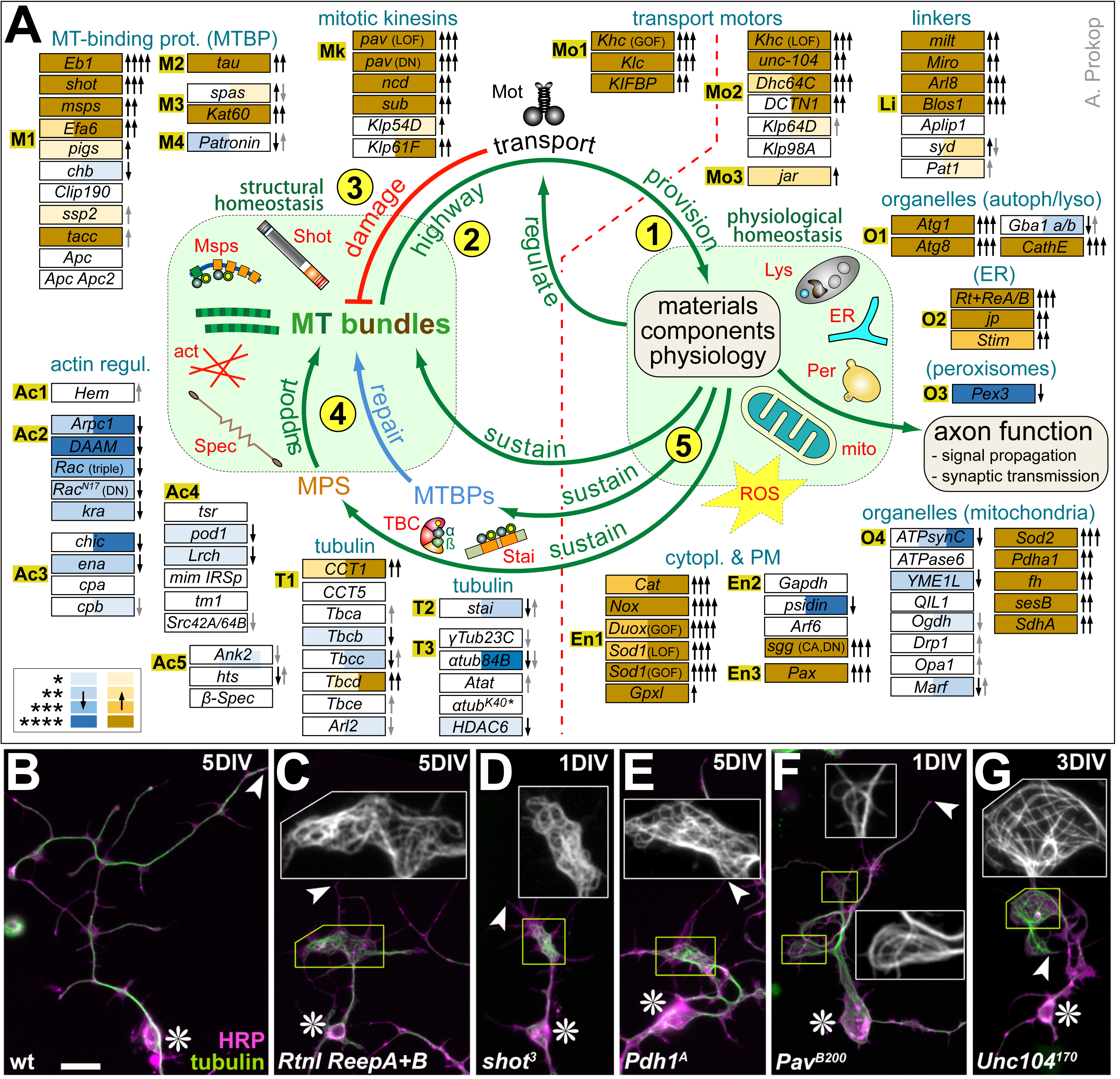
The dependency cycle of local axon homeostasis, the associated genetic loss-of-function conditions and their curling phenotypes. **A**) The centre depicts the dependency cycle of local axon homeostasis (Prokop, 2021a; Smith et al., 2023): (1) axonal transport provides materials and organelles to uphold axonal physiology and function; (2) MT bundles provide the highways for this transport; (3) axonal transport damages MT bundles and causes MT damage including curling; (4) MT-binding proteins (MTBPs) and the axonal cortex are required to maintain MT bundles, thus counterbalancing transport-induced damage; (5) transport-dependent components and physiology are required to uphold MT maintenance machinery. Some components are illustrated around the cycle: act, actin; ER, endoplasmic reticulum; Lys, lysosome; mito, mitochondrion; Mot, motor protein; Msps, Mini spindles (a MT polymerase); Per, peroxisome; ROS, reactive oxygen species; Shot, Short stop (actin-MT linker); Spec, spectrin (cortex component); TBC, tubulin-binding cofactor. Boxes on the outer ring list genetic loss-of-function conditions grouped by protein classes as indicated by orange-highlighted code: ‘Ac1’, Scar/WAVE complex; ‘Ac2’, actin nucleation regulators; ‘Ac3’, actin plus end-binding; ‘Ac4’, other actin regulators; ‘Ac5’, cortical actin-regulating or -binding; ‘En1’, ROS-regulating; ‘En2’, glycolysis component; ‘En3’, integrin adhesion components; ‘Li’, axonal transport cargo linkers; ‘M1’, MT plus end-binding; ‘M2’, MT lattice-binding; ‘M3’, MT severing; ‘M4’, MT minus end-binding; ‘Mk’, mitotic kinesins; ‘Mo1’, mutations causing hyperactivation of kinesin-1 or -3; ‘Mo2’, MT-binding transport motors (LOF); ‘Mo3’, actin-binding minus-end directed motor (myosin 6); ‘O1’, (auto-)phago/lysosome-regulating; ‘O2’, ER-regulating; ‘O3’, peroxisome regulating; ‘O4’, mitochondrial factors; ‘T1’, tubulin chaperones; ‘T2’, α/β-tubulin heterodimer-binding; ‘T3’, tubulins and tubulin acetylase. The dashed line proposes which classes of genes affect either structural or physiological homeostasis when functionally deficient. Black or grey arrows indicate the degree of MT-curling increase (pointing up) or decrease (pointing down; see Tab.S1 for values); colours match those in Tab.S1 and indicate statistical significance as indicated bottom left (* P ≤ 0.05; ** P ≤ 0.01; *** P ≤ 0.001; **** P ≤ 0.0001; see values in Tab.S1). **B-G**) Examples of MT curling in *Drosophila* primary neurons. Images show neurons of different genotypes (bottom left; see Tab.S1) cultured for different lengths of time (top right; DIV, days in vitro) and stained for tubulin (green) and the neuronal surface marker HRP (magenta). Cell bodies are indicated by asterisks, axons tips with arrowheads. Yellow boxes demarcate areas of MT-curling and are shown as 2.5-fold magnified greyscale insets (tubulin channel only). Scale bar in A represents 20µm in all images. Raw data files are available on figshare.com (https://doi.org/10.6084/m9.figshare.33112271; Tab.S1 folder).

**Fig. 2.**
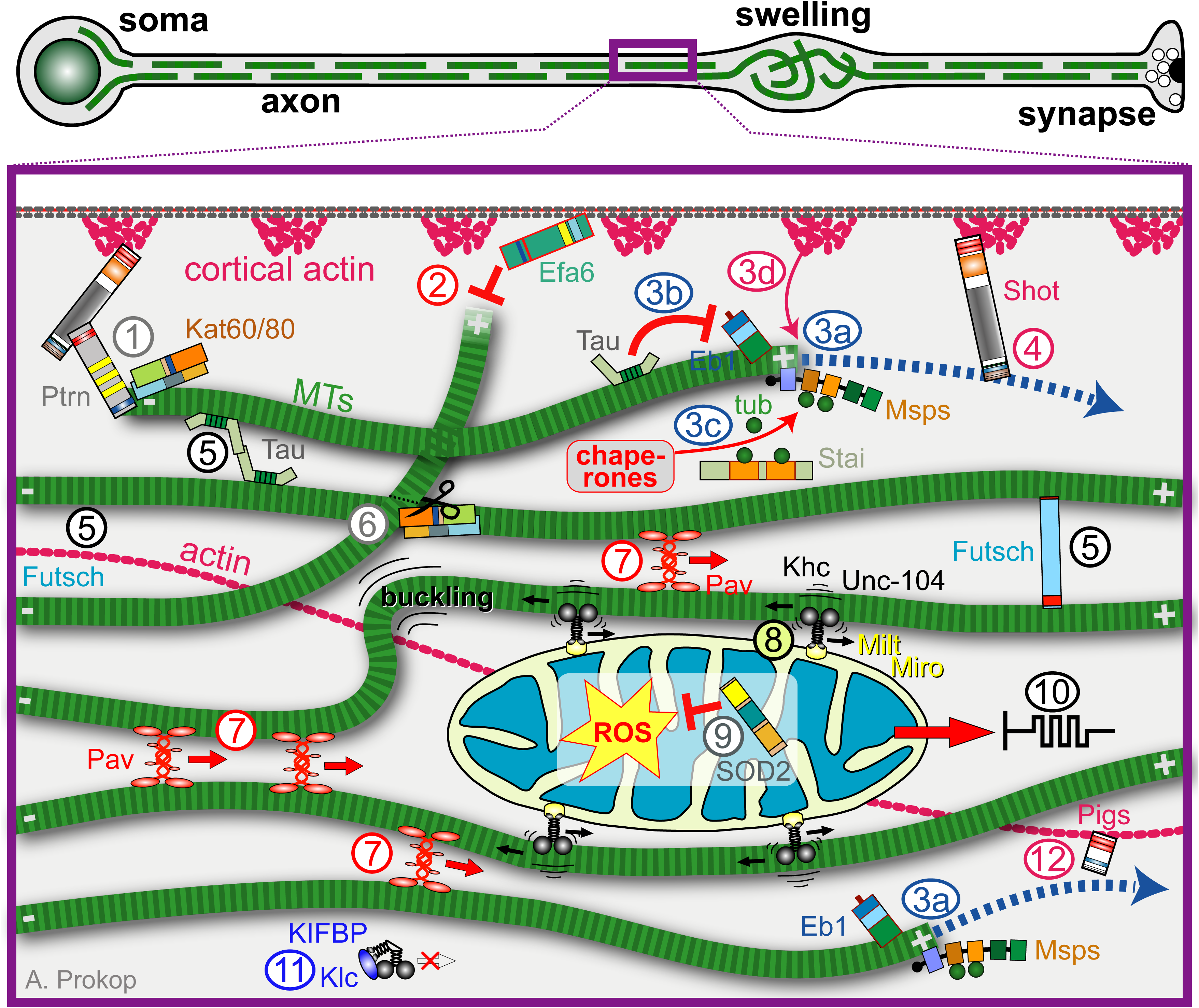
Illustration summarising published and proposed molecular mechanisms relating to MT-curling. The image on top illustrates a neuron with a continuous MT bundle (light/dark-green-dashed lines) and a pathological swelling with curled MTs. The close-up below shows molecular mechanisms discussed in the main text: **(1)** hypothesised MT minus-end anchorage to the axonal cortex; **(2)** Efa6-mediated cortical inhibition of MT polymerisation of MTs that have left the parallel MT bundles and project towards the plasma membrane; **(3)** the machinery maintaining MT polymerisation: (3a) the scaffold protein Eb1 and MT polymerase Msps mutually maintain their localisation at MT plus ends upholding MT polymerisation, (3b) Tau prevents Eb1 binding to MT lattices preventing its plus-end depletion, (3c) tubulin biogenesis feeds MT polymerisation processes, (3d) cortical actin rings promote MT polymerisation through mechanisms still unknown; **(4)** Shot-Eb1-mediated guidance of MT extension along the cortex into parallel bundles; **(5)** Futsch-mediated MT-MT cross-linkage; **(6)** Katanin-mediated severing of MTs that cross each other; **(7)** MT zippering to prevent buckling; **(8)** anterograde transport (here of a mitochondrion) causing MT-buckling; **(9)** Sod2 as an example of an enzyme involved in maintaining ROS homeostasis; **(10)** cytoplasmic resistance to cargo transport; **(11)** cargo-free transport motors detach from MTs in an inactive λ configuration stabilised by factors such as Klc or KIFBP; **(12)** hypothetical Pigs-mediated guidance of MT extension along central actin filaments.

Here we put the dependency cycle model to the test. We demonstrate that axonal MT-curling is initiated in healthy and mutant neurons alike, but only wild-type neurons maintain structural homeostasis by re-establishing parallel bundles. Our genetic manipulations of 105 genes from a broad cell biological spectrum reveal that ∼40% of manipulated genes can cause the common MT-curling phenotype. Closer analyses of 20 curl-inducing conditions consistently support the predicted binary divide in a structural and a physiological homeostasis component of the circle. Our data support the dependency cycle’s validity suggesting it as a useful concept for wider application in the neurodegeneration field.

## Results

### Live imaging reveals reduced reversibility of MT-curling events in mutant neurons

Disorganised MT-curling in axons is a potential pathological indicator not only in cultured *Drosophila* neurons, but it has likewise been observed *in vivo* in the fly brain (Hahn et al., 2021; Okenve-Ramos et al., 2024; Qu et al., 2019) and in axons of vertebrate and mammalian neurons (Hahn et al., 2019; Prokop, 2021a; Smith et al., 2023; see Discussion). To gain a better understanding of disorganised MT-curling we studied the origins and timeline of curling events. For this, we carried out live-imaging experiments using either laser scanning or spinning disc microscopy of primary neurons. We used wild-type neurons and neurons functionally deficient for either of three genes known to cause severe MT-curling in fixed preparations (Tab.S1): (1) the actin-MT crosslinking factor Short stop (allele used in homozygosis: *shot^3^*; Sánchez-Soriano et al., 2009), (2) the MT plus end-associating factor Eb1 (*Eb1^04524^*; Hahn et al., 2021) and (3) the kinesin-1-associated protein Kinesin light chain (*Klc^8ex94^*). Shot and Eb1 are both involved in MT bundle maintenance (Qu et al., 2022; orange-highlighted ‘M1’ in Fig.1A and ‘1’, ‘3a’ and ‘4’ in Fig.2), whereas Klc is required for the inactivation of the Kinesin heavy chain motor protein (‘11’ in Fig.2) with its functional loss causing motor hyperactivation (Liew et al., 2026).

In our first attempts, MTs were labelled with the live-dye SiR-tubulin (Lukinavičius et al., 2014). Since SiR-tubulin is a taxane derivative expected to convert GDP-tubulin into a GTP-like expanded and stabilised state (Siahaan et al., 2022), we suspected it might partially suppress MT-curling. We therefore also tried Jupiter^Ubi.mCherry^ (a fluorescently tagged MT-associated protein expressed by a ubiquitin promoter; Karpova et al., 2006). The Jupiter^Ubi.mCherry^-labelled neurons consistently showed more severe MT-curling than observed with SiR-tubulin (compare ‘SiRtub’ and ‘Jup’ for the same genotypes in Fig.3D).

**Fig 3.**
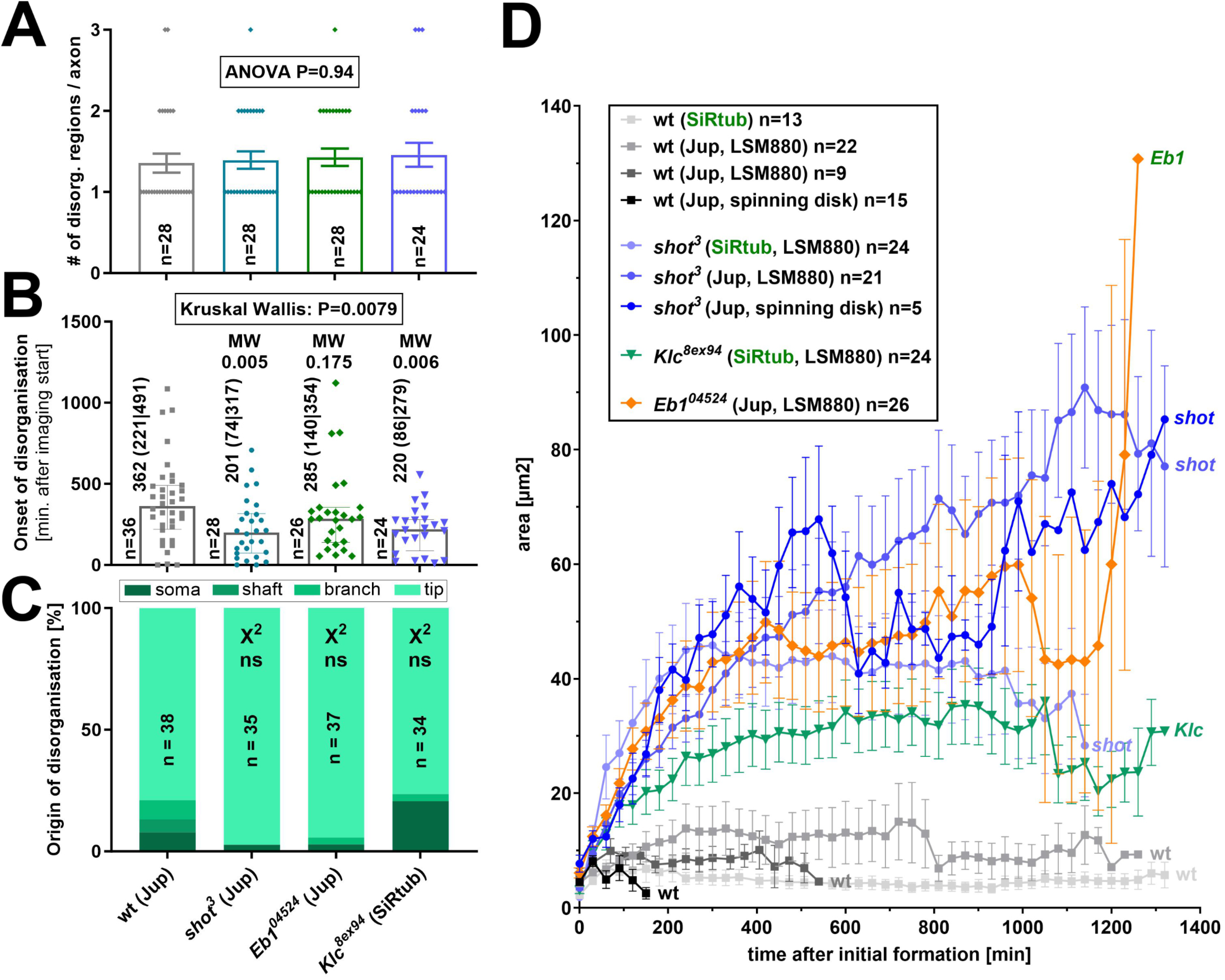
Analysis of origins and expansion of areas of disorganisation in axons. (**A**) Number of disorganised regions per axon. (**B**) Onset of axonal MT-curling relative to the imaging start. (**C**) Origin of disorganised regions in neurons, classified into soma, axon shaft, branch and tip. (**D**) Development of the total area of MT-curling along axons from initiation (set as ‘0’) to the end of the observation period; neurons from different genetic conditions (wt, wild-type; *Eb1*, *Eb1^04524/04524^*; shot, *shot^3/3^*; *Klc*, *Klc^8ex94,8ex94^*) were labelled either with SiR-tubulin (SiRtub) or *Jupiter^Ubi.mCherry^* (Jup) and imaged using a laser scanning (LSM880) or spinning disk microscope, as indicated and colour coded in the inset which also provides sample numbers of neurons imaged per condition. Raw data files are available on figshare.com (https://doi.org/10.6084/m9.figshare.33112271).

In all examples, live imaging started 2 hrs after plating of the neurons and continued for up to 22 hrs (Fig.3D). During the entire imaging period, wild-type and mutant neurons alike displayed on average ∼1.3 curl-initiation events along their axons (Fig.3). The median time when MT-curling first occurred differed only moderately between genotypes ranging from 201 to 362 min across genotypes, with wild-type having a potential bias for later and *shot^3^* and *Klc^8ex94^*for earlier onset; cases of very late onset were the exception (∼1,100 minutes in wild-type and *Eb1^04524^* mutant neurons, and ∼500 to 700 minutes in *shot^3^* and *Klc^8ex94^*mutant neurons; Fig.3B). MT-curling events were initiated (i) close to the soma, (ii) along the axon shaft, (iii) at axonal branch points, or (iv) in growth cones at axon tips (Figs. 3C, S1; Mov.S1). These areas might provide the necessary space for individual MTs to curl which would otherwise be prevented by plasma membrane rigidity. In agreement with this hypothesis, most curling events for all four genotypes initiated in growth cones at axon tips which are usually the most spacious axon compartments (Figs. 3C, S1E,F).

Whereas these parameters appeared rather similar across genotypes, clear distinctions were observed when monitoring the dynamic changes of individual MT-curling areas from the timepoint of their initiation to the end of the documentation period (Fig.3D): in wild-type neurons, areas of curling were clearly contained (Mov.S1) whereas mutant neurons, especially those labelled with Jupiter^Ubi.mCherry^, caused a steady increase over time with only transient periods of mild recovery (Fig.3D).

Taken together, our findings of comparable MT-curling initiation frequencies with preferential onset in growth cones across genotypes - but persistence of MT-curling only in mutant conditions - are consistent with the homeostasis model (see Discussion).

### Large-scale studies reveal MT-curling in ∼40% of manipulated genes

To identify potential mechanisms causing MT-curl induction, we assessed MT-curling phenotypes of a wide range of mutant conditions, using the MT disorganisation index (MDI; see Methods) as quantifiable readout. Here we present data from experiments that cover 200 different conditions and/or manipulations for 105 different genes (Tab.S1; Fig.1). About half of these data were from unpublished experiments (‘new’ in the last column of Tab.S1) or the re-analysis of images from published experiments not previously analysed to the common standards applied here (‘re’). The remaining data were collated from published work already using the standards (references provided in last column of Tab.S1). *Drosophila* primary neurons were usually generated from whole embryos which were homozygous mutant or displayed gene knock-down in all neurons (‘e’ in Tab.S1;Prokop et al., 2012; Voelzmann and Sánchez-Soriano, 2022). In some cases, phenotypes may be transiently masked by maternal contribution, which is persisting wild-type product deposited in the eggs by heterozygous mothers (Roote and Prokop, 2013). Such maternal contribution may persist for several days but can be overcome through different strategies: (i) by culturing cells for several days *in vitro* (indicated as ‘eDIV’ in Tab.S1), (ii) by pre-culturing them in a centrifuge tube for about 5 days before plating and growing them for usually a day (‘pre’ in Tab.S1; Prokop et al., 2012), or (iii) by extracting neurons from late larval brains if homozygous individuals survive to that stage (‘Li’ in Tab.S1).

Of the 105 studied genes (most validated with independent genetic tools; Tab.S1), an astonishing ∼40% displayed MT curling (light/dark-brown highlighted in Fig.1A and Tab.S1). To spot rules or patterns in this unprecedented data set, we grouped genes by function. They comprise tubulins and tubulin chaperones or regulators (orange-highlighted ‘T1-T3’ in Fig.1A), actin regulators (‘Ac1-Ac4’), MT-binding proteins (MTBPs; ‘M1-M4’), mitotic kinesins (‘Mk’), transport motor proteins (‘Mo1-3’; see legend for details), cargo linkers (‘Li’), genes regulating organelles or their function (‘O1-4’), as well as cytoplasmic enzymes (‘En1,2’). When aligning these classification with their appropriate position around the dependency cycle (Fig.1A), it becomes clear that dysfunction of very different aspects of axonal cell biology can cause the shared MT-curling phenotype (Fig.1B-G). This is in clear agreement with our hypothesis that the dependency cycle can be broken in various position to cause a shared or common disease outcome (see Introduction). In the following we will explain the data we obtained for the various gene groups.

### Different gene groups trigger MT-curling through distinct mechanisms

Nine of the assessed gene groups have prominent links to MT-curling (‘M1’, ‘Mk’, ‘Mo1’, ‘Mo2’, ‘L’, ‘O1’, ‘O2’, right column of ‘O4’, ‘En1’ in Fig.1A). For some of these, the mechanisms through which their malfunction causes curling were described previously, but will be briefly summarised here.

Within the ‘M1’ and ‘M2’ classes, the MT plus-end associated proteins Eb1, Msps and Shot together with the MT lattice binder Tau, were shown to guide the extension of proliferating MT plus ends into parallel bundles (‘3a/b, 4’ in Fig.2; Alves-Silva et al., 2012; Hahn et al., 2021). Efa6 is a cortical collapse factor eliminating extending MTs that escape guidance and leave the parallel bundles, thus providing quality control (‘2’ in Fig.2; Qu et al., 2019). Loss of any of these factors would be expected to affect structural homeostasis by reducing bundle maintenance (‘4’ in Fig.1A), hence cause MT-curling without ROS involvement. We tested this for Shot by treating *shot^3^*mutant neurons (displaying prominent MT-curling; Fig.1D) with the vitamin E analogue Trolox (6-hydroxy-2,5,7,8-tetramethylchroman-2-carboxylic acid). Trolox is known to act as a beneficial antioxidant in many cell systems (including neuronal models of neurodegeneration) by inhibiting fatty acid peroxidation and quenching singlet oxygen and superoxide (Chow et al., 1994; Giordano et al., 2020; Janc and Müller, 2014). Trolox failed to suppress MT-curling in *shot^3^* mutant neurons (Fig.4A), thus agreeing with our prediction that ROS is not involved in this phenotype.

**Fig. 4.**
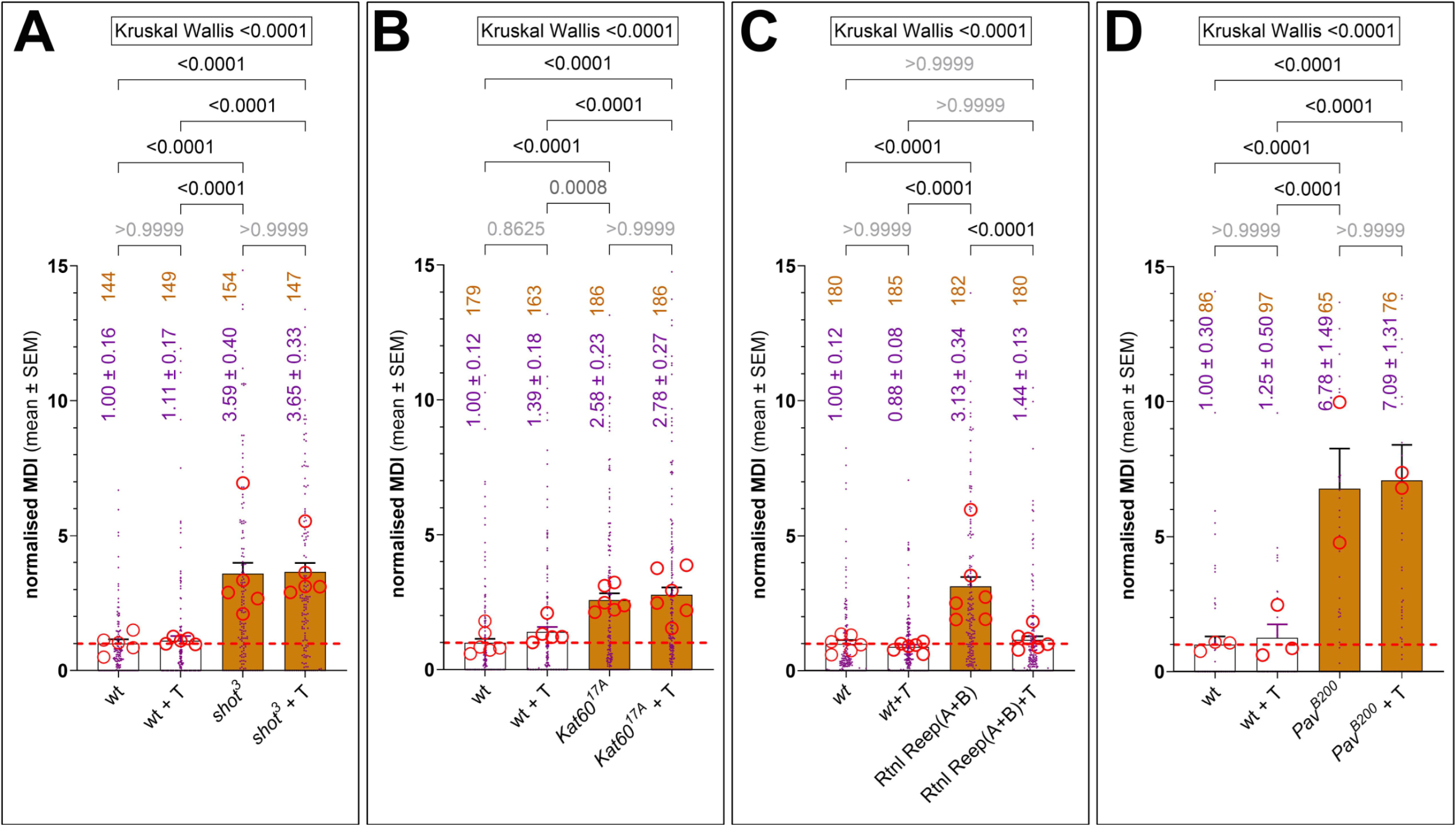
New experiments assessing MT-curling phenotypes with Trolox. Each graph represents one set of experiments composed of wild-type without Trolox (wt), wt with Trolox (+T) and the respective loss-of-function mutant condition (as indicated in A-D) without/with Trolox. MT-curling is measured as MT disorganisation index (MDI; area of MT curling divided by the product of axon length and an assumed axons width of 0.5µm), with bars representing mean ± standard error of mean all normalised to their wild-type controls without Trolox (stippled red line); blue dots reflect data for individual neurons, the red circles the means of repeats (usually 20-30 neurons per individual coverslip). Blue numbers indicate the mean ± SEM of pooled data, the orange number the total number of neurons analysed, and Dunn’s multiple comparison results are indicated above bars in black (significant) or grey (not significant); the intensity of brown fill-colour of bars reflects the degree of statistical significance relative to wild-type control without Trolox. Raw data files are available on figshare.com (https://doi.org/10.6084/m9.figshare.33112271).

The ‘Mo1’ group comprises mutant conditions that cause hyperactivation of transport motors, thus causing increased transport activity; this is expected to overwhelm the MT maintenance machinery, hence derail structural homeostasis and cause ROS-independent MT-curling. Consistent with our model, MT-curling in all tested conditions (*Klc^8ex94^*, *Khc^E177K,R947E^*, KIFBP knock-down, targeted expression of KIF5A^Δex27^) was not rescued by Trolox (Liew et al., 2026; Fig.5).

**Fig. 5.**
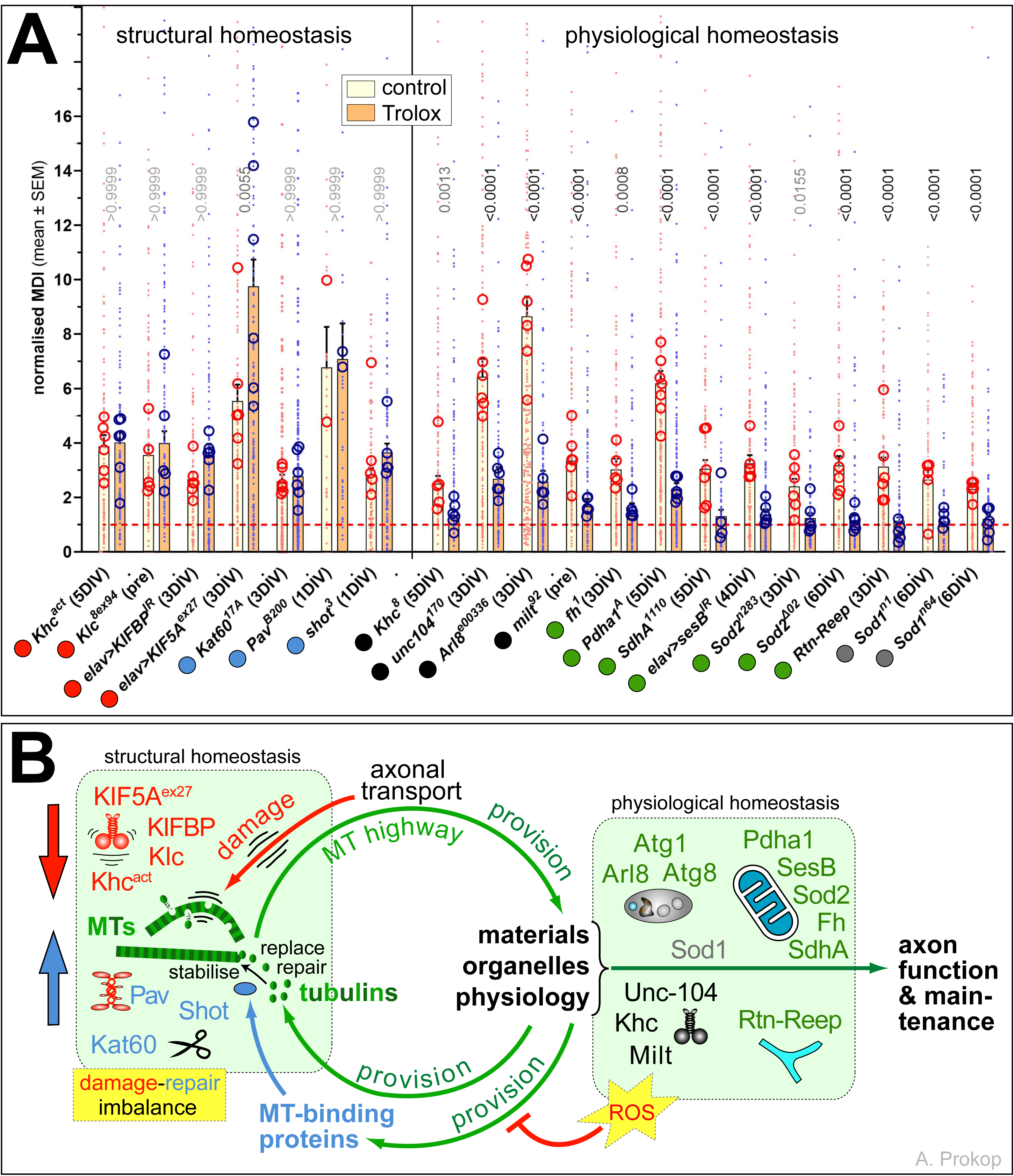
Summary of all experiments to date using Trolox applications. **A**) Graph comparing the MT-curling phenotypes of neurons carrying certain loss-of-function mutant conditions (light brown) normalised to untreated control neurons (red stippled horizontal line) and compared to the same mutant condition but treated with 100 nM Trolox applied with the culture medium (darker brown); genetic manipulations on the left side of the vertical line represent either functional loss of MT-binding proteins (indicated by blue dots) or hyperactivation of transport motors (red dots), none of which is rescued by Trolox. Manipulations on the right side cause MT-curling rescued by Trolox; they comprise conditions of transport deficiency (black dots), organelle dysfunction (green dots) or the cytoplasmic ROS regulator Sod1 (grey dot). Most data were published previously (*Khc*, *Kif5A*, *KFBP*, *Klc*, *Arl8*, *milt*, *unc-104* in Liew et al., 2026; *fh*, *Pdha*, *SdhA*, *sesB*, *Sod2* in Murray-Cors et al., 2025; *Sod1* in Shields et al., 2025); details of new data (*kat60*, *Pav*, *shot*, *Rtn-Reep*) are show in Fig.4. **B)** Results from A presented in the context of the ‘dependency cycle of local axon homeostasis’ (see Fig.1). <u>Light green box on the left</u>: it contains all tested genetic manipulations frm the left side of the stippled separator line in A (font colours corresponding to colour of dots in A) supporting the ROS-independent damage-repair imbalance hypothesis (red and blue arrows) referred to as ‘structural homeostasis’ (above box). <u>Light green box</u> <u>on the right</u>: it lists all tested genetic manipulations from the right side of the stippled separator line in A, consistently supporting the hypothesis that derailing ‘physiological homeostasis’ causes ROS-dependent MT-curling; note that information about the autophagosome regulators Atg1 and Atg8 is taken from a recent preprint (Gupta et al., 2026). Raw data files are available on figshare.com (https://doi.org/10.6084/m9.figshare.33112271).

In contrast, the ‘Mo2’ and ‘Li’ groups contain mutant conditions causing MT-curling by abolishing functions of transport motors or of those linkers that mediate transport of mitochondria or endo/lyso/autophagosomes (Fig.1G; ‘8’ in Fig.2). Here we further complemented this gene group by showing that MT-curling is triggered also by functional loss of Blos1 (*Blos1^ex2^*), an obligatory member of the BLOC1 complex mediating Arl8 recruitment and endo/lysosomal vesicle transport (Boda et al., 2019; Lund et al., 2021; Fig.1A and Tab.S1). Of the tested genes in these groups (loss of transport motors Khc and Unc-104 and the linker complex members Milt and Arl8) all cause ROS-dependent MT-curling consistent with physiological dyshomeostasis (Liew et al., 2026; Fig.5).

Beyond transport deficits of organelles, MT-curling is also observed when disturbing physiological homeostasis directly, as previously demonstrated for functional loss of the five proteins mediating mitochondrial physiology (right column of ‘O4’; Fig.1,E; Murray-Cors et al., 2025), direct manipulations of ROS-regulating enzymes (‘En1’; Liew et al., 2026; Shields et al., 2025), or loss of the autophagosome regulators Atg1 and Atg8 (‘O1’; Gupta et al., 2026). Here we show that also functional loss of factors relating to endoplasmic reticulum (ER) morphogenesis or physiology (‘O2’) cause MT-curling, as revealed by neurons triple-deficient for the three morphogenetic proteins Rtnl1, ReepA and ReepB (Yalçin et al., 2017) or lacking functions of the ER contact-site components Jp/JPH1-3 or Stim (Fig.1A,C and Tab.S1). Whether manipulations of lysosome physiology likewise trigger MT-curling remains to be seen; first analyses of lysosomal enzymes (‘O1’ in Fig.1) show no effect when deleting glucocerebrosidase (Gba1a/b) but strong MT-curling when knocking down the predicted orthologue (CG5863) of the aspartic endopeptidase Cathepsin E (Tab.S1, Fig.1A). For all tested factors from these groups (Sod1, Sod2, Pdha1, SesB/ANT, Fh, SdhA, Atg1, Atg8, and the triple combination of Rtnl1, ReepA and ReepB), MT-curling was consistently ROS-dependent (Figs.4C, 5; Gupta et al., 2026; Murray-Cors et al., 2025; Shields et al., 2025).

Our analyses of the ‘M3’ group of MT-severing AAA ATPases (McNally and Roll-Mecak, 2018) revealed prominent MT-curling upon loss of the catalytic Kat60 subunit of Katanin. This finding might relate to Katanin’s ability to sever MTs which cross each other (McNally and Roll-Mecak, 2018; Vemu et al., 2018; Wang et al., 2017), thus acting as a control factor removing non-aligned MTs in areas of curling (‘6’ in Fig.2). Alternatively, it may interact with Patronin/CAMSAP (Jiang et al., 2018) to repair sites of MT damage (Zhang et al., 2023) or mediate anchorage and length regulation of MT minus ends (‘1’ in Fig.2; Akhmanova, 2018; Akhmanova and Steinmetz, 2015; Dong et al., 2017; Nashchekin et al., 2016; Ning et al., 2016; Noordstra et al., 2016). Any of these functions would be expected to sustain structural homeostasis, and this agrees with our finding that Kat60 loss causes ROS-independent MT-curling (Figs.4B, 5). Loss of Spastin, another MT-severing AAA ATPase, was reported to cause axonal swellings with curled MTs in mouse neurons (Fassier et al., 2013; Tarrade et al., 2006). In contrast, even stringent mutant conditions for Spastin in *Drosophila* primary neurons did not cause this phenotype and failed to enhance *Kat60^17A^*-mediated curling, as assessed in *Kat60^17A^ spas^575^* double-mutant neurons, suggesting that Spas and Kat60 are not functionally redundant in this context (Fig.S2A).

The ‘Mk’ group comprises mitotic kinesins, and we found MT-curling upon functional loss of at least three of them: the kinesin-6 family members Pav/KIF23 and Sub/KIF20A and the kinesin-14 family member Ncd/KIFC1 (Fig.1E; details in Fig.S3). Mitotic kinesins obtained their name from their role in sliding antiparallel MTs in the mitotic spindles but were later found to be present in parallel MT bundles of axons where they can cross-link MTs (Guha et al., 2021). For the mammalian mitotic kinesins KIF11, KIF15 and KIF1C it was shown that they can simultaneously move on two parallel MTs and bundle them *in vitro* (Drechsler and McAinsh, 2016; Kapitein et al., 2005; Oladipo et al., 2007; Tao et al., 2006). We hypothesise therefore that mitotic kinesins might zip MTs together, thus stabilising bundles against MT buckling, especially behind large, transported cargoes (‘7’ in Fig.2; Bicek et al., 2009). This could work anterogradely (plus end-directed Pavarotti and Subito) and retrogradely (minus end-directed Ncd; Cai et al., 2009; She and Yang, 2017; Wendt et al., 2002). Such a function would contribute to structural homeostasis, which agrees with our finding that MT-curling upon Pavarotti loss is ROS-independent (Figs.4D and 5).

Taken together, factors from very different aspects of axonal cell biology can cause the common MT-curling phenotype, and for 20 tested factors the ROS-in/dependence of MT-curling is consistent with the binary homeostasis concept proposed by the dependency model.

### Loss of actin regulators tends to reduce MT-curling and increase axon length

Three groups of genes are less prone to cause MT-curling, as illustrated by the white and blue colour code in Fig.1A and Tab.S1. These include a group of mitochondrial factors (left column of ‘O4’ in Fig.1), actin regulators (‘Ac2-5’) and tubulin regulators (‘T1-3’).

The left column of ‘O4’ comprises six important regulators of mitochondrial fission, fusion and cristae morphogenesis which jointly suggest that mitochondrial contortion does not cause oxidative stress during the 5-day period these neurons were analysed (discussed in Murray-Cors et al., 2025).

‘Ac1-Ac5’ In Fig.1A lists the actin regulators we analysed. This group shows a clear trend to reduce MT-curling (Fig.6A). This might be explained through two distinct mechanisms: Firstly, loss of actin regulators often affects the integrity of cortical actin maturing into the membrane-associated periodic skeleton (Qu et al., 2017; Xu et al., 2013). Cortical actin loss negatively impacts MT polymerisation, expected to reduce the likelihood that MTs extend out of the bundle and initiate MT-curling (Qu et al., 2017). Secondly, loss of many actin regulators causes growth cones to be narrow (Gonçalves-Pimentel et al., 2011; Qu et al., 2022; Sánchez-Soriano et al., 2010), thus limiting the space required to initiate MT-curling (as mentioned above; ‘tip’ in Fig.3C). Since narrow growth cones also force polymerising MTs to extend straight, i.e. in parallel to the axon axis, this can be expected to enhance axon growth (Prokop et al., 2013) and might explain the trend of longer axons we observed (Fig.6B). This said, extra growth due to narrow growth cones seems to be cancelled out if cortical actin is affected and MT polymerisation along axons reduced (see details in Fig.S4).

**Fig. 6.**
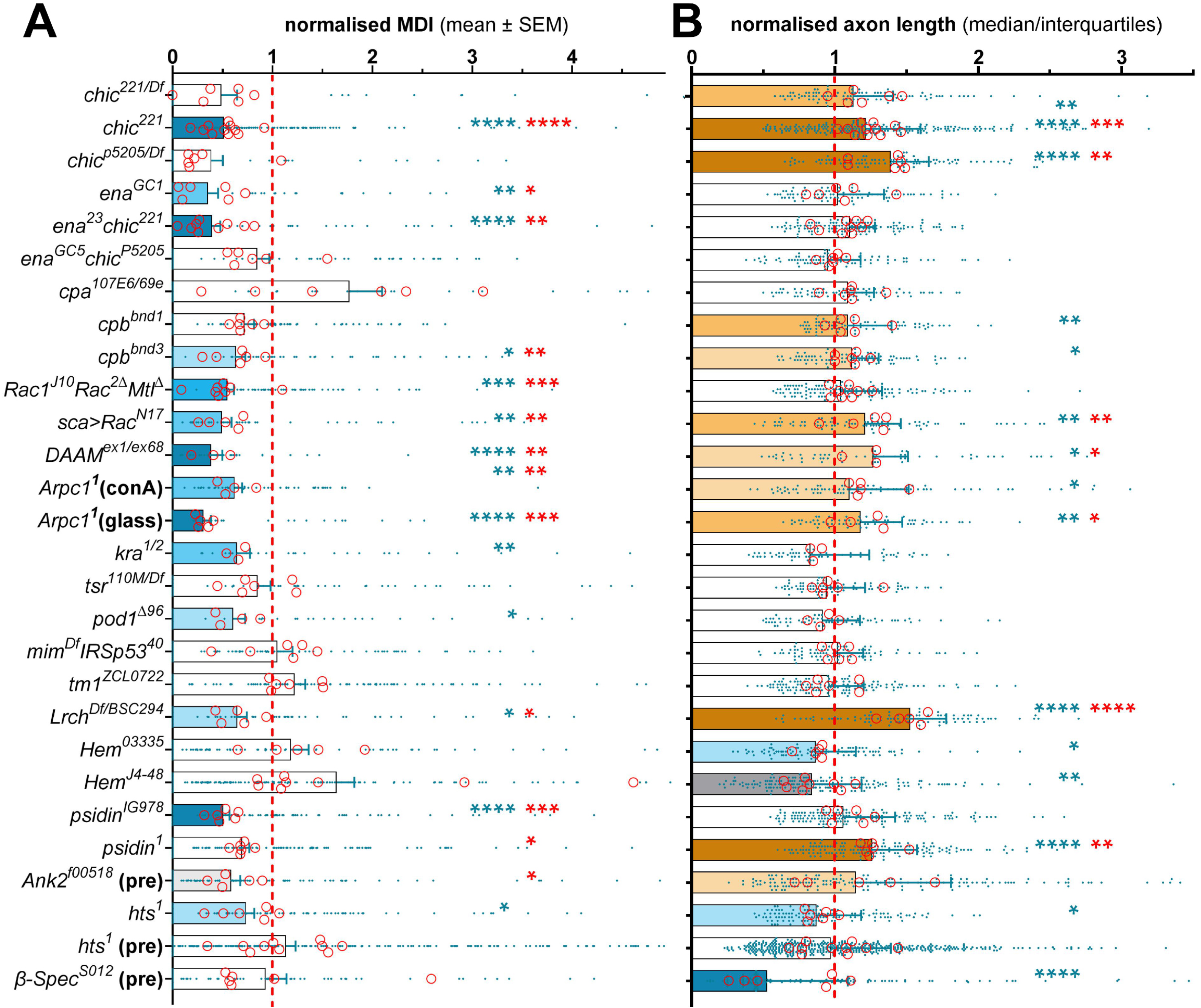
Loss of actin regulators shows a trend towards reduced MDI and enhanced axon growth. Analyses of different actin regulators; for detailed data and gene information see Tab.S1. Most of the mutant neurons carry strong loss-of-function mutant alleles either in homozygosis (no slash in superscript), over a deficiency uncovering the gene in question (allele/Df), or in heteroallelic combination with another strong loss-of-function mutant allele (allele/allele); note that three examples display double-mutant and one example a triple-mutant condition, that *sca>RacN17* represents the overexpression of a dominant-negative version of Rac1 in otherwise wild-type mutant background, and that Arpc1 analyses are shown for neurons grown on glass or using concanavalin-A as substrate (conA). (**A**) MT-curling is measured as MT disorganisation index (MDI; area of MT curling divided by the product of axon length and an assumed axons width of 0.5µm), with bars representing mean ± standard error of mean all normalised to parallel controls (stippled red line); blue dots reflect data for individual neurons, the red circles the means of repeats (usually 20-30 neurons per individual coverslip). Blue asterisks indicate the significance of pooled data established using Mann-Whitney tests, which is also reflected in the bar colour, the red asterisks show the significance of repeat means established with t-tests. For bar colours see inset in Fig.1 and asterisks indicate the following values: * P ≤ 0.05; ** P ≤ 0.01; *** P ≤ 0.001; **** P ≤ 0.0001 (see exact values in Tab.S1). (**B**) Graphs and colour codes for axon length measurements follow the same logic, but bars reflect the median and upper interquartile range. Raw data files are available on figshare.com (https://doi.org/10.6084/m9.figshare.33112271; see in Tab.S1 folder).

The strongest axon elongation was observed upon loss of the actin-binding factor Lrch (Fig.6B). This phenotype is rescued by targeted expression of GFP::Lrch which localised throughout axons (without obvious F-actin association; Fig.S5). So far, Lrch deficiency in flies was shown to be a viable condition with shortened lifespan that causes female sterility and affects cell division in S2 cells (Foussard et al., 2010). LRCH proteins uniquely combine actin-binding calponin-homology domains with leucin-rich repeats and some of its four human paralogues have little understood links to diseases (Rivière et al., 2020), where *Drosophila* might provide the means to explore the underlying mechanisms.

Taken together, losses of actin-binding regulators in *Drosophila* primary neurons tend to supress MT-curling, which is frequently correlated with increased axon growth in vitro; potential explanations were provided. Our findings agree with the fact that mutations of human actin regulators mostly relate to neurodevelopmental rather than neurodegenerative diseases (Suppl. Mat. in Smith et al., 2023).

### Losses of tubulins or tubulin chaperones cause reduced axon growth but no MT-curling

The ‘T1-T3’ group of genes shown in Fig.1A also has little tendency to cause MT-curling. It comprises the MT nucleation factor γTub23C/TUBG1,2 and various regulators involved in tubulin provision, in particular the α1-tubulin-encoding gene *αtub84B/TUBA* and members of the CCT and TBC chaperone complexes essential for tubulin biogenesis (‘3c’ in Fig.2; Pinho-Correia and Prokop, 2023; Voelzmann et al., 2016). Losses of most of these factors lead to shorter axons (Fig.7) likely caused by reduced MT polymerisation (less net MT extension) or MT nucleation (lower MT numbers; Hahn et al., 2021; Roostalu et al., 2020), expected to reduce the likelihood of MT-curling as discussed in the previous section. Our studies further suggest that tubulin biogenesis mediated by these factors might, in part, take place locally in axons of *Drosophila* primary neurons (Fig.S6A-Q). But this is likely negligible and irrelevant in the short neurites of primary *Drosophila* neurons where axonal transport is not a limiting factor (Fig.S6R) – but could be relevant in longer vertebrate axons.

**Fig. 7.**
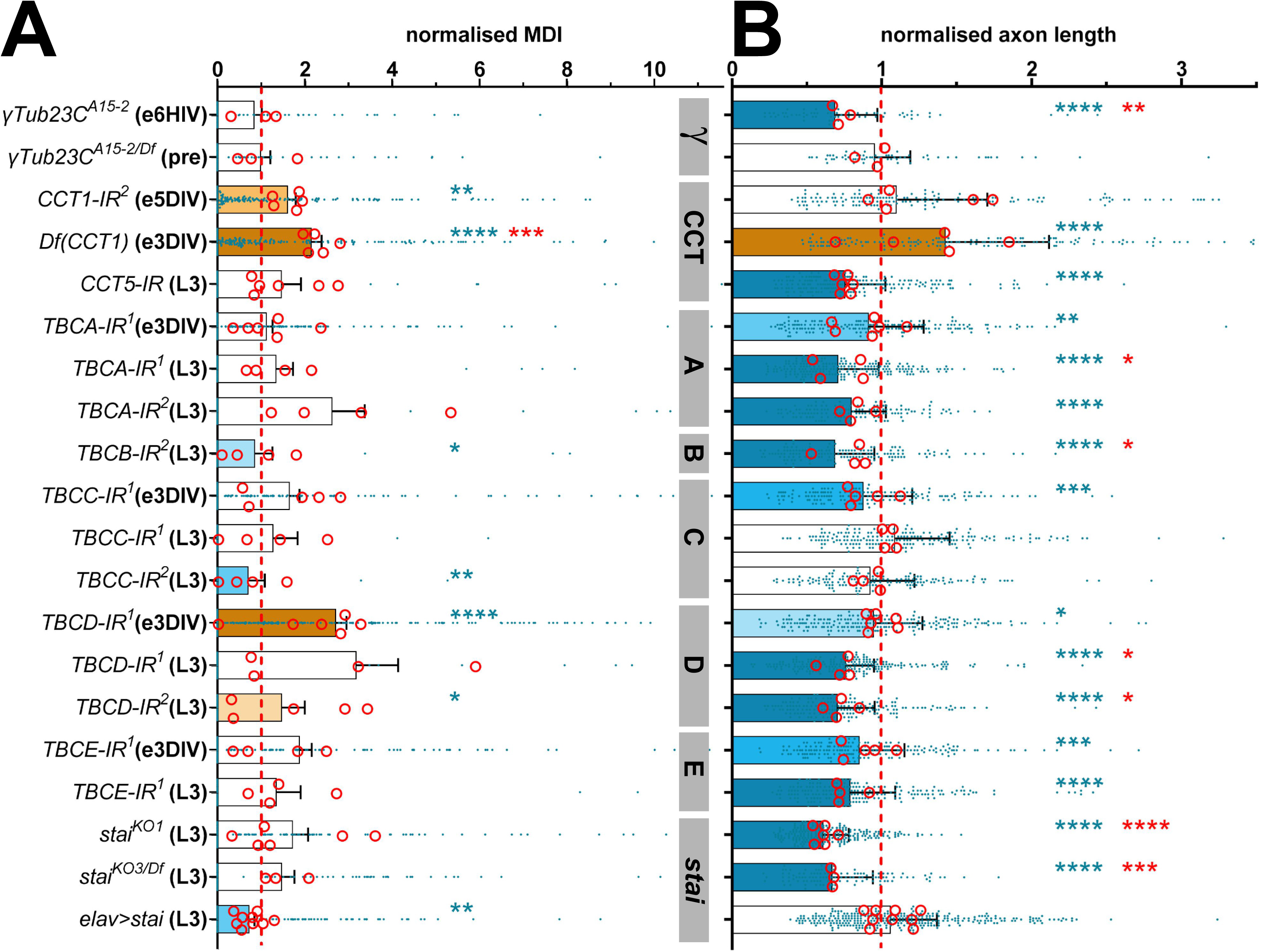
Analysis of chaperones involved in tubulin biogenesis. Analyses of the normalised MDI (**A**) and normalised axon length (**B**) presented with the same logic as explained in Fig. 6 (for detailed data see Tab.S1). The suffix ‘-IR’ (interference RNA) in gene names refers to gene knock-down driven by *elav-Gal4*, with TBCA-D being knocked down with two independent transgenic constructs (indicated by superscript ‘1’ and ‘2’); further conditions represent *elav-Gal4*-driven overexpression of *stai* or loss-of-function mutant alleles either in homozygosis (no slash in superscript) or over deficiency uncovering the respective gene (allel/Df). To facilitate reading of this graph, grey boxes between A and B indicate the different genes manipulated and listed below B. Cultured neurons were either embryo-derived and cultured for 3 days (e3DIV) or derived from larval brains and cultured for 1 day (L3). Raw data files are available on figshare.com (https://doi.org/10.6084/m9.figshare.33112271; see in Tab.S1 folder).

Of the tested tubulin-linked chaperones (‘T1’ in Fig.1A), only CCT1 and potentially TBCD cause MT-curling (Fig.7A). This might be explained through chaperone-independent functions of these proteins (Pinho-Correia and Prokop, 2023). For example, curl-inhibiting functions of CCT1 might be explained by its ApiCCT1 domain originally discovered in the CCT1-orthologue of the yeast *Saccharomyces* (Fig.S7). Yeast ApiCCT1 was shown to suppress pathological protein aggregates and revert neuronal atrophy in mouse Huntington’s disease models (Sontag et al., 2013; Tam et al., 2006; Zhao et al., 2016).

Comparable to tubulin chaperone deficiencies, also loss of Stathmin (Stai/STMN1-4; ‘T2’ in Fig.1A) does not induce MT-curling but reduces axon growth (Fig.7), as was similarly observed upon loss of the mammalian orthologue STMN2 (aka SCG10; Manna et al., 2007). This phenotype might likewise be caused through loss of tubulin leading to reduced MT polymerisation. This was suggested by our quantitative qRT-PCR analyses of wild-type and Stai-deficient larval CNSs (details in Fig.S8), which agree with previous reports of reduced tubulin protein levels extracted from whole *stai* mutant specimens (Duncan et al., 2013). We propose that Stai binds and protects tubulin pools from auto-inhibitory regulation of its own biogenesis (‘3c’ in Fig.2; Pinho-Correia and Prokop, 2023; Sellin et al., 2008; Voelzmann et al., 2016) - although tubulin-independent roles of Stathmin during axon growth cannot be excluded (Beccari et al., 2025).

Taken together, functional deficiencies of tubulin, tubulin-chaperones, γ-tubulin, stathmin and MPS components work through very different mechanisms, but are all expected to share reduced MT nucleation/polymerisation as a common denominator that could explain their trend for reduced MT-curling.

### Acetylation of α-tubulin’s K40 has no obvious impact on MT-curling

The final group of genes we tested are the enzymes Atat and HDAC6 involved in acetylation and deacetylation of α-tubulin’s lysin 40 (α^K40^; located inside the MT lumen; Fig.8A; Hubbert et al., 2002; Luo et al., 2025) which we assigned to the ‘T3’ group of tubulin regulators. We focussed on these enzymes because α^K40^ acetylation is easy to detect (6-11-B1 antibody specific for α^K40^ acetylation; Fig.8B; Piperno and Fuller, 1985) and is known to be increased in response to various cellular stressors, often taken as a readout for MT stability (Donker and Godinho, 2025). α^K40^ acetylation was shown to render MTs more flexible and less prone to breakage (Eshun-Wilson et al., 2019; Li and Yang, 2015; Portran et al., 2017; Xu et al., 2017), and we wondered whether it might therefore influence curling behaviour. However, when using the complete loss-of-function mutant alleles *Atat^15^* and *HDAC6^KO^* (viable in homozygosis) we did not observe obvious MT-curling phenotypes in primary neurons (Fig.8G; Tab.S1), although functional loss of these enzymes was confirmed by changes in the observed acetylation patterns (Figs.8C and S9C).

**Fig. 8.**
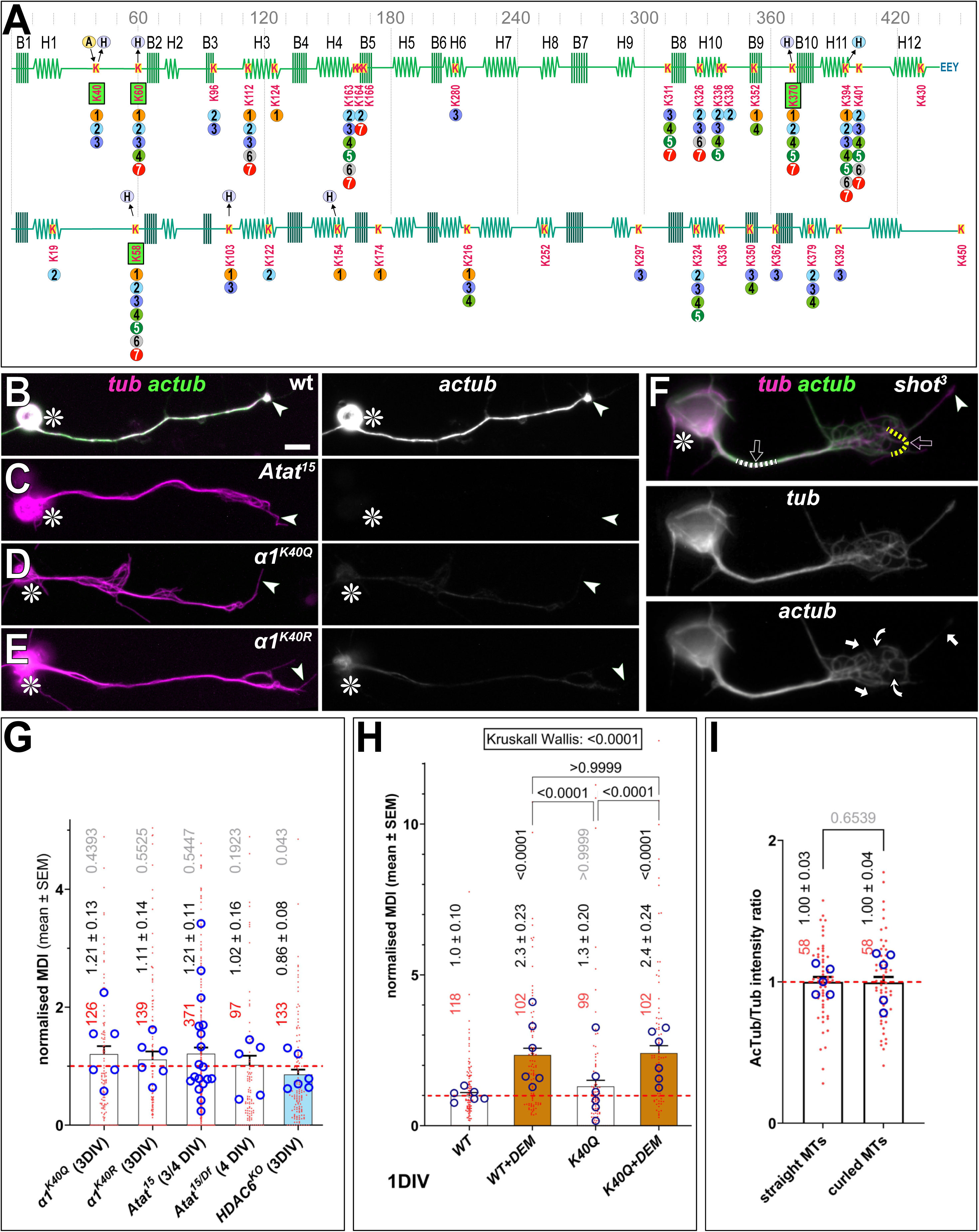
MT curling is not influenced by tubulin acetylation. **E**) The secondary structure of α- (light green) and ß-tubulin (dark green) showing the positions of ß-sheets (B1-10) and α-helices (H1-12; image modified from Keskin et al., 2002); the scale above indicates residue numbers. All lysins are shown (K) with their position indicated below (luminal lysins are green emboxed). ‘A’ in the orange circle indicates Atat acetylase activity, ‘H’ in blue circles the deacetylase activity of HDAC6. Coloured circles below lysins indicate reports of acetylation on specific lysine residues from independent mass spectrometry studies; numbers in circles refer to the following studies: [1] material obtained from wild-type and HDAC6-deficient mouse brains (Liu et al., 2015a); [2] from 16 different rat tissues (Lundby et al., 2012); [3] from HeLa cells (Liu et al., 2015b); [4] from HeLa cells (Hansen et al., 2019); [5] from mouse embryonic fibroblasts (Weinert et al., 2018); [6] from *Drosophila* (Weinert et al., 2011); [7] from MV4-11, Jurkat and A549 cells (Choudhary et al., 2009). **B-E**) Primary *Drosophila* neurons (asterisks, cell bodies; arrowheads, axon tips) double-stained with a general anti-tubulin antibody (magenta) and the 6-11-B1 antibody specific for K40-acetylated α-tubulin (green, and shown in grey scale on the right). In Atat-deficient neurons, 6-11-B1 staining is completely absent (A,A’), in neurons homozygous for the two K40 mutant alleles staining is almost completely abolished (C-D’). **F**) A *shot^3^* mutant neuron stained as in B-E, indicating segmented lines tracing MTs in bundled (white stippled line) and curled (yellow stippled line) areas used for intensity measurements, curved arrows indicate MTs in curled areas lacking acetylation and the straight arrow an MT lacking distal acetylation. The scale bar in B represents 10 µm in B to E and 5 µm in F. **G-I**) Quantification of MT-curling in different mutant neurons (G), in wild-type and α1^K40Q^ neurons with and without DEM treatment (H) and of acetylated tubulin to total tubulin intensity ratios in bundled (white stippled line in F) versus curled (yellow stippled line) areas; graphs constructed as explained in Fig.4. Raw data files are available on figshare.com (https://doi.org/10.6084/m9.figshare.33112271).

To further validate these findings, we tested neurons homozygous for the viable mutant alleles *α1^K40Q^* and *α1^K40R^* which mimic the acetylated and non-acetylated state of α1, respectively (Jenkins et al., 2017). This approach is particularly powerful in *Drosophila* neurons which, almost exclusively, express only one α- and one β-tubulin isotype (α1 aka αTub84B and β1 aka βTub56D; details in Fig.S8E). Therefore, modifications of these tubulins in these neurons are expected to be fully penetrant and undisturbed by redundancies with other tubulin isotypes. This notion is corroborated by our finding that *α1^K40Q^* and *α1^K40R^* homozygous mutant neurons show almost no immunoreactivity for acetylated α^K40^ (Fig.8D,E). Despite their obvious mutant state, these mutant neurons displayed no obvious changes in MT-curling at 3 DIV (Fig.8G), thus agreeing with our findings for Atat1 and HDAC6 and with reports that they display very mild neuronal phenotypes in *Drosophila in vivo* (Jenkins et al., 2017). To challenge this further, we induced MT-curling independently by applying the ROS-elevating compound diethyl maleate (DEM) to *α1^K40Q^*mutant neurons, but found no obvious trends to suppress or enhance the phenotype (Fig.8H).

*Vice versa*, we assessed whether the curling of MTs has an impact on α1 acetylation. We first used the strongly curling *shot^3^* mutant neurons and measured the ratio of staining intensities of acetylated versus total tubulin obtained from MTs that were either in bundled or in curled areas; we did not observe obvious differences between these regions (Fig.8F,I). We then used curl-inducing conditions to assess whether patterns of MT acetylation were affected in curled areas. This approach was incentivised by reports in HeLa cells where damage through the hyperactivated transport motor variant *KIF5A^Δex27^::GFP* (a strong-curl-inducing condition in our neurons: Liew et al., 2026) caused entire MTs to lose their α^K40^ acetylation, mediated by the HDAC6 deacetylase (Andreu-Carbó et al., 2024). We therefore analysed neurons that expressed *KIF5A^Δex27^::GFP* or were homozygous for *HDAC6^KO^*, as well as neurons that were wild-type or homozygous for *shot^3^* and *Khc^E177R+R947E^* (another hyperactivated transport motor condition). In HDAC6-deficient neurons, acetylated and total tubulin staining tended to be completely overlapping across the entire MT network including curled areas and filopodial tips (Fig.S9C). In all other conditions, including wild-type and *KIF5A^Δex27^::GFP* expression, a small fraction of neurons displayed a moderate subset of non-acetylated MTs in curled areas (curved arrows in Figs.8F and S9), only MTs extending into filopodia often lacked acetylation of their distal segments (white arrows in Figs.8F and S9). It appears therefore that MTs in *Drosophila* neurons are better protected from deacetylation than in HeLa cells, which might be due to potential higher abundance of lattice-binding factors such as Tau or Ensconsin/MAP7 (Andreu-Carbó et al., 2024).

Taken together, our data suggest that α^K40^ acetylation and MT-curling are not interdependent processes in these neurons. Since ∼20 tubulin lysins were shown to be acetylated (of which four, including α^K40^, are located inside the MT lumen; green highlighted in Fig.8A), α^K40^ acetylation might reflect a broader pattern of MT acetylation which, in sum, may have impact and influence MT-curling.

## Discussion

### Data consistently support the dependency cycle model

The dependency cycle of local axon homeostasis was proposed as a potential model providing explanations for why different forms of neurodegeneration, such as motoneuron disease, hereditary spastic paraplegia or Charcot-Marie-Tooth disease can all be caused by mutations in genes of very different functions (Fig.1A; Prokop, 2021a; Smith et al., 2023). To put the model to the test, we performed live imaging and reported data for 105 genes obtained from generating and analysing new or re-analysing old data sets, complemented by published data that match our experimental and statistical procedures. This represents an unprecedented data pool not only regarding the numbers covered but also the standardisation of their analyses in one normed primary neuron system, using MT-curling as an efficient and relevant readout. Through this standardisation, data can be compared and contrasted, hence classified, thus pinpointing common themes and trends where derived hypotheses can be tested with efficient *Drosophila* genetics.

We strongly feel that our data consistently support the ‘dependency cycle of local axon homeostasis’ model. Firstly, our live imaging data reveal similar frequencies and sites of MT-curling initiation for wild-type and mutant neurons, but curling is contained only in wild-type neurons. These observations agree with the ‘structural homeostasis’ concept which proposes a constant threat of MT-curl initiation through off-track polymerisation or motor transport-induced forces (‘3’ in Fig.1), kept in check by maintenance through MT-regulating proteins (‘4’ in Fig.1), but derailed if motors are hyperactive or MTBPs functionally impaired.

Secondly, our extensive manipulations of 105 genes clearly show that different groups of genes (as defined by their function; Fig.1A; Tab.S1) can cause one common MT-curling phenotype. This agrees with the proposed idea that the dependency cycle can be broken in different positions, mirroring the situation observed in neurodegeneration (see Introduction; Smith et al., 2023; Wilson III et al., 2023). Notably, there are clear trends for different gene groups as to whether they cause MT-curling, and potential mechanistic explanations for each gene group were addressed in the Results part.

Furthermore, our data agree with the predicted classification into two major gene groups (note the dashed separator line in Fig.1A): genes influencing MTs directly cause ROS-independent curling when functionally impaired; genes required to uphold cellular or organelle physiology cause ROS-dependent curling if deficient (summarised in Fig.5; further discussed below). This classification might bring further order into the complex genetic landscape that underpins neurodegeneration and could provide interesting food for thought when contemplating potential therapeutic strategies.

Taken together, we feel that the dependency model provides helpful ideas and concepts to address neurodegeneration, but it might also provide important explanations for the often-mentioned statement that ageing is the biggest risk factor for neurodegeneration: during ageing many aspects of neural biology are functionally weakened (Salvadores et al., 2017), and this is likely to increase the cycle’s vulnerability and lower its threshold to collapse upon genetic, acquired or environmental insults.

### MT-curling as a relevant readout and proxy for neurodegeneration

The key readout chosen here is disordered curling as a feature of axonal MT bundle decay. These bundles are key to axonal transport, thus providing the essential lifeline for axons. Their decay, as indicated by MT-curling, is therefore likely to turn them into the axons’ Archilles heels paving a path to degeneration. Based on this notion we propose MT-curling as a likely proxy for neurodegeneration, as is further supported by other observations: Firstly, MT-curling does not only occur in *Drosophila* neurons but is likewise found in vertebrate neurons upon similar genetic or pharmacological manipulations (Ahmad et al., 2006; Dalpe et al., 1998; Hahn et al., 2021; Liew et al., 2026; Patil et al., 2025; Rafiq et al., 2022; Sánchez-Soriano et al., 2009; Voelzmann et al., 2026). Secondly, MT-curling is found *in vivo* in the *Drosophila* brain, both upon ageing or when applying conditions known to induce curling in culture, including MTBP deficiencies, manipulations of autophagy or ROS dyshomeostasis (Gupta et al., 2026; Hahn et al., 2021; Okenve-Ramos et al., 2024; Qu et al., 2019; Shields et al., 2025). In the *Drosophila* brain, MT decay and curling has been clearly shown to be causative for other hallmarks of neuronal atrophy (Gupta et al., 2026; Okenve-Ramos et al., 2024; Shields et al., 2025), agreeing with the central role given to MTs in the dependency model. This said, MT curling may appear a less likely event *in vivo* when considering that initiation of MT-curling requires space in form of widened areas of axons (see our live imaging: Fig.3) and that axons in the brain are constricted by axonal fasciculation and glia wrapping. However, axonal blebbing has been reported *in vivo* in *Drosophila* (e.g. Okenve-Ramos et al., 2024), potentially providing the conditions where MT-curling can initiate. Similarly, also mammalian neurons *in vivo* show swellings with MT-curling, in all cases associated with decay due to neurodegenerative predisposition or ageing (Dalpe et al., 1998; Eyer et al., 1998; Fiala et al., 2007; Nag et al., 2020; Yang et al., 1999).

### Mechanisms underpinning ROS-dependent and -independent MT-curling

As proposed in previous work (Liew et al., 2026), ROS-independent MT-curling is triggered when disturbing the damage-repair balance underlying structural homeostasis. This occurs either upon motor hyperactivation or by weakening functional aspects of the MT maintenance machinery (‘1-7, 11,12’ in Fig.2) - comparable to potholes occurring on a highway upon excessive traffic or negligent highway maintenance (Shields et al., 2024). The ROS-independence of structural homeostasis has been previously demonstrated for motor hyperactivation, and here we now demonstrated that also the loss of MT regulators (Shot, Kat60, Pav) causes ROS-independent MT-curling (Figs.4A,B,D and 5).

For physiological dyshomeostasis, which occurs upon functional deficiency of organelle transport or of genes directly contributing to physiological processes, all tested examples consistently show ROS dependence of MT-curling (Fig.5). The potential mechanisms causing ROS dyshomeostasis through transport deficit, autophagosome or mitochondrial aberration were discussed elsewhere (Gupta et al., 2026; Liew et al., 2026; Murray-Cors et al., 2025). Here we added aberration of ER physiology as a further factor (Fig.6C), which likely relates to the ER’s role in calcium regulation, lipid metabolism, but also its contact sites with most other organelles and the plasma membrane - establishing means of intracellular coordination made possible by the continuous tubular ER networks pervading entire axons (Öztürk et al., 2020; Phillips and Voeltz, 2016; Smith et al., 2023).

Harmful ROS appears therefore the common denominator when derailing physiological homeostasis, but the mechanisms through which it triggers MT-curling remain less clear; this may involve the oxidation of MTs, of MT-binding proteins or of their respective upstream signalling molecules (see Discussions in Liew et al., 2026; Shields et al., 2025), and it may also involve ROS-induced blebbing of axons opening up the necessary space for curl induction (Mehta et al., 2025).

## Conclusions

In our view, the dependency cycle provides a powerful system in which to unravel new mechanisms or concepts, test hypotheses, and generate new ideas and understanding that can inform research into human neurodegeneration. One important limitation is the model’s restriction to neurons, not directly including those diseases that primarily affect glial cells (Gleichman and Carmichael, 2020; Verkhratsky et al., 2014). This said, any inherited, acquired or environmentally induced degenerative disorder of the nervous system will eventually render the pathology irreversible by affecting the neurons or their axons. Therefore, stabilising axons may be beneficial even upon non-autonomous insults.

Secondly, some may perceive the use of *Drosophila* neurons as a further limitation. However, *Drosophila* neurons provide the cost-effective and time-saving genetic strategies required to tackle complex axon biology. Such systematic work in *Drosophila* has laid foundations of enormous translational potential in the past leading to five (arguably 6) Nobel Prizes in “Physiology or Medicine” (Bellen et al., 2010; Prokop, 2018). As discussed above, there are indeed good indications already for the translational potential of the dependency model, but more data need to be gathered in mammalian or human studies. For this, it should be considered that immunohistochemical or ultrastructural work in higher organisms or humans often restricts to neurofilaments as readouts, thus sidelining MTs as the far more important cytoskeletal component and limiting our current subcellular insight into relevant pathological conditions (Hahn et al., 2019; Prokop, 2020).

## Acknowledgements

This work was made possible through funding by the Biotechnology and Biological Sciences Research Council through research grants to A.P. (BB/P020151/1, BB/L000717/1, BB/M007553/1, BB/I002448/1, BB/C515998/1) and N.S.S. (BB/R018960/1, BB/W016907/1) and DTP studentship to R.B. (BB/D526561/1), W.C. (BB/T008725/1) and L.P.C. (number not known). Further funding supporting this work was awarded by the Wellcome Trust to A.P.; (077748/Z/05/Z), the German Research Council (DFG) to A.V. (VO 2071/1-1), the Leverhulme Trust (ECF-2017-247) and Royal Society (RGS\R2\222151), BBSRC (UKRI1894/APP46392) and Academy of Medical Sciences (SBF008\1140) to I.H., FCT PhD scholarship from the Portuguese Foundation for Science and Technology Foundation to C.G.P., the University of Hull startup funding to A.V. and by parents to Y.-T.L. and Y.Q. The Fly Facility has been supported by funds from The University of Manchester (https://www.bmh.manchester.ac.uk/research/support/funding/strategic) and the Wellcome Trust (087742/Z/08/Z; AP). We are indebted for the continued support by the Fly Facilty manager Sanjai Patel and grateful for the services of the Bloomington Drosophila Stock Center (NIH P40OD018537) from which most of the used fly stocks were obtained, and for the services of the flybase.org and ensembl.org data bases which enormously facilitated the essential background searches and data mining exercises hat were so crucial for this work. The funders had no role in study design, data collection and analysis, decision to publish, or preparation of the manuscript.

## Methods

### Flies

To obtain information about suitable fly stocks, expression patterns, human orthologues, gene functions, the official gene names and the related literature, we used the latest versions of FlyBase (Jenkins et al., 2022; Öztürk-Çolak et al., 2024). The fly stocks we used are listed below in alphabetic order of their official gene names, and their references and sources are provided; if obtained from stock centres, ‘BDSC’ indicates the Bloomington *Drosophila* Stock Center (bdsc.indiana.edu), ‘Kyoto’ the KYOTO Stock Center (kyotofly.kit.jp), and ‘VDRC’ the Vienna *Drosophila* Resource Center (shop.vbc.ac.at).

- *Ank2^f00518^* (BDSC #85608; Thibault et al., 2004)
- *Apc^Q8^* (Ahmed et al., 1998; Akong et al., 2002)
- *Apc2^g10^ (Akong et al., 2002)*
- *Aplip1**^EK4^*** (BDSC #24632; Horiuchi et al., 2005)
- *Aplip1**^Df^*** (*Df(3L)Fpa2*; BDSC #24633)
- *Arf6^GX16[w-]^* (BDSC #60585; Huang et al., 2009)
- *Arl2^Δ156^* (Chen et al., 2016a; deletes -189 to +1551 relative to the Arl2 ATG start codon; courtesy H. Wang)
- *Arl2^Δ309^* (Chen et al., 2016a; deletes -185 to +399 relative to the Arl2 ATG start codon; courtesy H. Wang)
- *Arl8^e00336^* (*PBac(RB)Arl8e00336*; BDSC #17846; Rosa-Ferreira et al., 2018)
- *Arl8^Df^* (*Df(3R)D7*; BDSC #1898; Rosa-Ferreira et al., 2018)
- *Arpc1^1^* (aka *Sop2^1^* Hudson and Cooley, 2002)
- *Atat^15^* (BDSC #602674; Niu et al., 2023)
- *Atat^Df^* (Df(3L)BSC113; BDSC#8970)
- *Atg1^Δ3D^* (courtesy Samantha Loh)
- *Atg8a^KG07569^* (*P{SUPor-P}Atg8a^KG07569^*; BDSC#14639)
- *ATPsynC^KG01914^* (Lovero et al., 2018; *ATPsynC^KG01914^*; P-element insertion in non-coding 5’ exon generating a protein null; BDSC#13923)
- *ATPsynC^Df^* (Parks et al., 2004;*Df(3R)Exel6218*; uncovering ATPsynC; BDSC#7696)
- *Blos^ex2^* (courtesy Ole Kjaerulff; Cheli et al., 2009)
- *Cat^n1^* (Mackay and Bewley, 1989; Matthias Landgraf)
- *UAS-CathE^IR^* (*CG5863*; *P{TRiP.HMS05358}attP40*; BDSC#64022)
- *UAS-CCT1^IR#1^* (*P{GD10468}v34070/TM3;* VDRC#34070)
- *UAS-CCT1^IR#2^* (*P{y[+t7.7] v[+t1.8]=TRiP.HMS00639}attP2*; BDSC #32854; Perkins et al., 2015)
- *UAS-CCT5^IR^* (*P{KK108095}VIE-260B*; VDRC#109505)
- *Df(CCT1)* (Df(3R)Exel6191, P{w[+mC]=XP-U}Exel6191; BDSC #7670)
- *chb^2^* and *chb^4^* (Inoue et al., 2000; courtesy of D. van Vactor)
- *chb^Df^* [*Df(3L)BSC553*; BDSC #25116]
- *chic^221^* (Verheyen and Cooley, 1994; Wills et al., 1999; D. van Vactor)
- *chic^5205a^* (*P(Morrow et al.)chic05205a*; Wills et al., 1999)
- *chic^Df^* [*Df(2)Gpdh1A*; Wills et al., 1999]
- *Clip190^KO^* (Dix et al., 2013)
- *Clip190^Df^* (*Df(2L)CLIP-190* ; Beaven et al., 2015)
- *cpa^107E^* (Janody and Treisman, 2006)
- *cpa^69e^* (Gates et al., 2009; F. Janody)
- *cpb^bnd1^* (Delalle et al., 2005; F. Janody)
- *cpb^bnd3^* (Delalle et al., 2005; F. Janody)
- *DAAM^Ex68^* (Matusek et al., 2006; J. Mihaly)
- *DAAM^Ex1^* (Matusek et al., 2006; J. Mihaly)
- *DCTN1-p150/Glued^Δ22^* (Siller et al., 2005; removes entire locus and part of CG8833)
- *Dhc64C^4-19^* (Gepner et al., 1996; BDSC #5274)
- *Drp1^T26^* (Verstreken et al., 2005; BDSC #3662)
- *UAS-Duox* (Ha et al., 2005; Matthias Landgraf)
- *spas^5.75^* (Sherwood et al., 2004)
- *Eb1^04524^* (Elliott et al., 2005; Spradling et al., 1999)
- *Eb1^Df^* (*Df(2R)Exel6050*; BDSC #7532)
- *Eb1^5^* and *Eb1^04524^* (Elliott et al., 2005)
- *Efa6^GX6[w-]^* (Huang et al., 2009)
- *Efa6^GX6[w+]^* (Huang et al., 2009)
- *Efa6^Df1^* (*Df(3R)Exel6273*; BDSC #7740)
- *Efa6^Df2^* (*Df(3R)ED6091*; BDSC #9092)
- *Efa6^IR^* (*P{GD14945}v42321*; VDRC #42321)
- *elav-Gal4* (BDSC #8765)
- *ena^23^* (Ahern-Djamali et al., 1998; B. Baum)
- *ena^GC1^* (Ahern-Djamali et al., 1999; BDSC #8569)
- *ena^GC5^* (Gertler et al., 1995; BDSC #8570)
- *fh^1^* (Chen et al., 2016b; *fh^1^*; lethal S136R point mutation; BDSC#67161)
- *UAS-fh^IR^* (Anderson et al., 2005; *P{UAS-fh.RNAi.A}2*; BDSC#24620)
- *UAS-Gapdh^IR^* (*Gapdh1^GD7467^*; VDRC #31631)
- *Gba1a+b^GBA1ΔTT^* [*Df(3R)GBA1ΔTT*; BDSC#95249]
- *Gba1a+b^Exel6195^* [Df(3R)Exel6195; BDSC#7674]
- *UAS-*Gpxl^IR^ (BDSC#56992)
- *HDAC6^KO^* (BDSC#51182)
- *Hem^03335^* (hypomorphic allele; Schenck et al., 2004)
- *Hem^J4-48^* (amorph, W490term amino acid replacement; Hummel et al., 2000)
- *hts^1^* (Yue and Spradling, 1992)
- *IRSp53^40^* (Goddard, 2013; courtesy T. Millard)
- *jar^322^* (BDSC #8776; Morrison and Miller, 2008)
- *jar^Df^* (*Df(3R)Crb87-5*; BDSC #2363)
- *Jupiter^Ubi.mCherry^* (tagged MAP driven by ubiquitin promoter; Karpova et al., 2006; Lu et al., 2013)
- *UAS-jp^IR^* (VDRC ID #48530)
- *Kat60^17A^* (BDSC #64116; Mao et al., 2014)
- *Khc^8^* (Saxton et al., 1991; BDSC #1607)
- *Khc**^27^*** (Saxton et al., 1991)
- *Khc^Df^* (*Df(2R)BSC309*; Cook et al., 2012; BDSC #23692)
- *UAS-Khc^IR^* (*P{y[+t7.7] v[+t1.8]=TRiP.HMS01519}*; BDSC#35770)
- *Khc^E177K^* (Kelliher et al., 2018)
- *Khc^E177K,R947E^* (Kelliher et al., 2018)
- *UAS-KIFBP^IR^* (CG14043; BDSC #51858; Perkins et al., 2015)
- *Klc**^Df^*** (BDSC #7596)
- *Klc**^8ex94^*** (Gindhart et al., 1998; BDSC #31997)
- *Klp54D^IR^* (*Klp54D^HMJ30099^*; flybase.org: FBrf0208670; Ni et al., 2009; BDSC #63533)
- *Klp61F^urc-1^* (Wilson et al., 1997; BDSC #35508)
- *Klp61F^Df^* [*Df(3L)BSC311*; flybase.org: FBrf0200284; BDSC #24337]
- *Klp64D^k1^* (Ray et al., 1999; hypomorphic allele; BDSC #5578)
- *Klp64D^n123^* (Perez and Steller, 1996; BDSC #5674)
- *Klp98A^Δ47^* (Derivery et al., 2015)
- *kra^1^* (Lee et al., 2007)
- *kra^2^* (Lee et al., 2007)
- *Lrch^Df^* [*Df(2L)Lrch^Df^*; Foussard et al., 2010]
- *Lrch^BSC294^* [*Df(2L)BSC294*; 36C2-C9; Dix et al., 2013]
- *UAS-GFP-Lrch* (gift from François Payre; unpublished line generated in the context Foussard et al., 2010)
- *Marf^B^* (Sandoval et al., 2014; Hugo Bellen)
- *me31B-GFP* [*P{PTT-GB}me31B^CB05282^*; Buszczak et al., 2007; Weil et al., 2012]
- *mim^Df^* [*Df(2R)Exel6051*)](; Tom Millard)
- *milt**^92^*** (Cox and Spradling, 2006; Stowers et al., 2002)
- *milt**^Df^*** [*Df(2L)ED440, P{w[+mW.Scer\FRT.hs3]=3’.RS5+3.3’}ED440*; Ryder et al., 2004; Kyoto #150498]
- *Miro**^B682^*** (Guo et al., 2005; BDSC #52003)
- *Miro**^Sd32^*** (Guo et al., 2005; BDSC #95252)
- *msps^A^* (Hahn et al., 2021)
- *msps^146^* (Brittle and Ohkura, 2005)
- *mt:ATPase6^1^* (Celotto et al., 2006; *mt:ATPase6^1^*; lethal G116E point mutation)
- *Mtl^Δ^* (Hakeda-Suzuki et al., 2002)
- *ncd^1^* (Endow and Komma, 1997; Yamamoto et al., 1989; BDSC#1290; X-ray-induced 2.6kb deletion removing 5’ section of *ncd* extending *claret* locus)
- *UAS-Nox* (Liew et al., 2021)
- *Ogdh1^MI06026^* (Yoon et al., 2017; *Ogdh^MI06026-TG4.1^*; BDSC#77497)
- *Ogdh1^Df^* (Ryder et al., 2007; *Df(3L)Exel7253*; BDSC#7938)
- *Opa1^s3475^* (*P{lacW}Opa1^s3475^*; Spradling et al., 1999; BDSC #12188)
- *UAS-Opa^IR^* (*P{y[+t7.7] v[+t1.8]=TRiP.HMS00349}*; BDSC#32358; Perkins et al., 2015)
- *Pat1^robin^* (Loiseau et al., 2010)
- *Pat1^grive^* (Loiseau et al., 2010)
- *Patronin^c9-c5^* (Nashchekin et al., 2016)
- *Patronin^Df^* (*Df(2R)BSC355;* BDSC #24379; Cook et al., 2012)
- *pav**^B200^*** (BDSC #4384; Adams et al., 1998; Salzberg et al., 1994)
- *Pav^Df^* (*Df(3L)Exel9000*; BDSC #7921; Parks et al., 2004)
- *UAS-pav^DEAD^* (*UASp-GFP-pav.DEAD^37^*; Minestrini et al., 2002)
- *ubi-GFP-pav* (*Ubi-p63E-GFP-pav^53^*; Minestrini et al., 2002)
- *Pax^Δ1^* (deletes almost all of the Pax coding region, the downstream gene *CG31798*, and one of two *CG17544* transcripts; Bataillé et al., 2010)
- *Pdha1^A^* (Yamamoto et al., 2014; *Pdha^A^*, lethal G126E point mutation)
- *UAS-Pdha1^IR^* (Perkins et al., 2015; *P{TRiP.HMC04032}*; BDSC#55345)
- *Pex3^2^* (Faust et al., 2014)
- *UAS-Pfdn2-Venus* [*P{UAS-Pfdn2.Venus*; provided by Hongyan Wang;){Zhang, 2016 #9732}(]
- *pigs^1^* (Pines et al., 2010)
- *pod1^Δ96^* (Rothenberg et al., 2003)
- *psidin^1^* (Brennan et al., 2007; Stephan et al., 2012)
- *psidin^IG978^* (Stephan et al., 2012)
- *UAS-QIL1^IR^* (Guarani et al., 2015; Perkins et al., 2015; *P{TRiP.GLC01383}*; expression reduced to about 25%; loss of cristae junctions ; BDSC#44364)
- *Rac1^j10^* (Ng et al., 2002)
- *Rac2^Δ^* (Ng et al., 2002)
- *UAS-Rac^N17^* (Luo et al., 1994)
- *Rtnl1^1.w^* (O’Sullivan et al., 2012; Yalçin et al., 2017)
- *Rtnl1-YFP* (*PBac{681.P.FSVS-1}Rtnl1CPTI001291*; Cahir O’Kane; O’Sullivan et al., 2012)
- *ReepA^541^* (Yalçin et al., 2017)
- *ReepB^48^* (Yalçin et al., 2017)
- *UAS-Rpl10-GFP* [*P{w[+mC]=UAS-GFP-RpL10Ab}BF2b*; BDSC #42681]
- *sca-Gal4* (Mlodzik et al., 1990)
- *SdhA^1110^* (Mast et al., 2008; lethal E288K point mutation; BDSC#51659)
- *SdhA^1404^* (Mast et al., 2008; lethal V445E point mutation; BDSC#81120)
- *UAS-sesB^IR#1^* (Perkins et al., 2015; *P{TRiP.HMS01549};* BDSC#36661)
- *UAS-sesB^IR#2^* (P{TRiP.JF01528}; BDSC#31077; Perkins et al., 2015)
- UAS-sgg^S9A^ (constitutively active Shaggy/GSK3β; Hazelett et al., 1998)
- UAS-sgg^A81T^ (inactive form of Shaggy/GSK3β (Bourouis, 2002)
- *shot^3^* (Kolodziej et al., 1995)
- *Sod1^n1^* (Phillips et al., 1989; BDSC #24492)
- *Sod1^n64^* (Phillips et al., 1995; BDSC #7451)
- *Sod2^n283^* (Duttaroy et al., 2003; BDSC #34060)
- *Sod2^Δ02^* (flybase.org FBrf0208127; BDSC #27643)
- *UAS-Sod1* (J. Hu and J.P. Phillips, unpublished)
- *UAS-spas^IR^* (Trotta et al., 2004)
- *UAS-spas^K467R^* (Orso et al., 2005)
- *β-Spec^S012^* (C538T nucleotide substitution; Hülsmeier et al., 2007)
- *Src42A^E1^* (Tateno et al., 2000)
- *Src64B^ko^* (O’Reilly et al., 2006)
- *ssp2^ΔB^* (Gluszek et al., 2015)
- *stai^KO1^* (Yang et al., 2012; courtesy P. Rørth)
- *stai^KO3^* (Yang et al., 2012; courtesy P. Rørth)
- *stai^Df^* (*Df(2L)Exel6015*; Duncan et al., 2013)
- *UAS-stai* (UAS-insertion; courtesy P.Rørth)
- *UAS-Stim^IR^* (VDRC ID #47074)
- *sub^1^* (Jang et al., 2005; Schüpbach and Wieschaus, 1989; BDSC#5117; Q230term amino acid replacement)
- *syd**^Df^*** (*Df(3L)syd^A2^*; Bowman et al., 2000; deleting C-terminus; BDSC #32017)
- *syd**^z4^*** (Bowman et al., 2000; BDSC #32016)
- *tacc^1^* (Gergely et al., 2000)
- *tau^KO^* (Burnouf et al., 2016)
- *UAS-Tbca^IR1^* (*P{TRiP.HMJ21731}attP40*; BDSC #53677)
- *UAS-Tbca^IR2^* (*P{GD8629}v39893*; VDRC#39893)
- *UAS-Tbcb^IR2^* (*P{GD7246}v38466*; VDRC#38466)
- *ubi-Tbcb-GFP* [*P{Ubi-Tbcb.EGFP}*; provided by Antoine Guichet; Baffet et al., 2012]
- *UAS*-*Tbcc^IR1^* (*P{TRiP.HMC06477}attP40*; BDSC #67371)
- *UAS-Tbcc^IR2^* (*P{GD14747}v29360*; VDRC#v29360)
- *UAS*-*Tbcd^IR1^* (*P{TRiP.HMC06045*}*attP40*; BDSC #65170)
- *UAS-Tbcd^IR2^* (*P{GD12185}v27931*; VDRC#27931)
- *UAS*-*Tbce^IR1^* (*P{TRiP.HMS02865}attP40*; BDSC #44569)
- *UAS-Tbce-GFP* [*P{UASp-Tbce.EGFP}*; courtesy Giet Régis; Métivier et al., 2021]
- *Tm1^ZCL0722^* (Kim et al., 2011)
- *tsr^100M^* (Janody and Treisman, 2006)
- *tsr^Df^* [*Df(2R)or-BR6*; BDSC #36569)
- *αtub84B^Df^* (Df(3R)BSC633; BDSC #25724)
- *αtub84B+D^Df^* (*Df(3R)Antp17*; Duncan and Kaufman, 1975; Jenkins et al., 2017)
- *αtub84B^K40Q^* (BDSC #602672; Jenkins et al., 2017)
- *αtub84B^K40R^* (BDSC # 602673; Jenkins et al., 2017)
- *αtub84B^Df^* (*Df(3R)Antp17*; removing both *αtub84B* and *αtub84D*; Duncan and Kaufman, 1975)
- *γTub23C^A15-2^* (courtesy Paul Conduit; Vázquez et al., 2008)
- *γTub23C^Df^* [Df(2L)ED206; courtesy Paul Conduit]
- *tubP-Gal4* (courtesy L. Luo; Lee and Luo, 1999)
- *unc-104^170^* (Pack-Chung et al., 2007)
- *YME1L^del^* (Qi et al., 2016*YME1L^del^*; 2kb deletion removing most of the coding region; BDSC#95273)

Green balancer chromosomes used to identify mutant or construct-expressing embryos were readily available *FM7*, *CyO or TM3* balancers carrying *twi-Gal4 or Kr-Gal4* in combination with *UAS-GFP* or carrying a *Df-GFP* fusion construct (Casso et al., 2000; Halfon et al., 2002; Le et al., 2006).

### Cell culture and immunohistochemistry

#### Embryonic cultures

*Drosophila melanogaster* primary neuronal cell cultures were prepared as described previously (Prokop et al., 2012; Voelzmann and Sánchez-Soriano, 2022). Primary neurons were obtained either from embryos (stage 11 or older; stages according to Campos-Ortega and Hartenstein, 1997) or late larval CNSs. In brief, embryos were dechorionated with 50% bleach for 90 seconds, selected for the correct developmental stage or for genotype using green balancer chromosomes. Selected embryos were sterilised with 70% ethanol, washed in sterile-filtered growth medium (commercial Schneider’s medium, 20% fetal bovine, 2 mg/ml of human insulin, 1:1000 PenStrep) then mechanically dispersed with a pestle in dispersion medium followed by 10 min incubation at 37°C to allow for chemical dispersion (2ml 1.5xHBSS solution containing 100µg phenylthiourea, 30µl PenStrep, 2mg Collagenase, 4 mg Dispase II). After incubation, 200μl of Schneider’s medium was added, cells centrifuged at 750 rpm for 4 min, final volume of growth medium (30μl per culture chamber) was added to the pellet, cells suspended again, added to the chambers and closed with a Concanavalin A-coated coverslip sealed with vaseline. After 2hrs culturing coverslip-down at 26°C (for cells to settle and attach), chambers were turned upside-down (hanging-drop culture) for the further culture period.

#### Larval cultures

Late larvae were sterilised in ethanol, CNS dissected out in a drop of sterile Schneider’s medium in a sterile well, transferred to a centrifuge tube, the medium replaced by 200µl dispersion medium, mechanically dispersed with a pestle, then incubated for 10 min at 37°C. All following steps as for embryos.

#### Pre-cultures

For pre-cultures, procedures were followed as described for embryonic cultures up to the mechanical dispersion and centrifugation steps. Add 200 μl growth medium to the pellet, and incubate closed tube covered with aluminium foil in vertical position at 26°C for 3-7 days. After that, tubes were centrifuged, dispersion medium added to the pellet, solution carefully pipetted up and down, incubated at 37°C for 3-4 minutes, 200µl growth medium was added, tubes centrifuged again and then final volume of growth medium added to the pellet, dispersed and distributed to chambers.

### Antisera, smFISH and immunocytochemistry

Neuronal cultures were fixed over 30 mins at room temperature (RT) in 0.1 M phosphate buffer (pH 7.2) containing 4% paraformaldehyde and 0.05% glutaraldehyde. Staining was performed either in 0.2% Triton in PBS, or in PBS following 20 min incubation with 0.5% Tergitol in PBS. Antibody staining and washes were performed with phosphate buffered saline with 0.3% TritonX-100 (PBT). We used primary antibodies against the following antigens: α-Tubulin (rat, YOL1/34, 1:500 or DM1A, mouse, 1:1000), acetylated α-tubulin (6-11-B1, mouse, 1:1000; Piperno and Fuller, 1985), GFP (ab6556; rabbit; 1:1000), anti-puromycin (mouse; 1:1000; Kerafast), anti-dTBCB (1:500; courtesy Antoine Guichet; Baffet et al., 2012). Furthermore, we used fluorescently-conjugated anti-HRP (goat; 1:50; Jackson ImmunoResearch) and fluorescently conjugated AffiniPure® secondary antibodies (donkey; 1:400).

### Puromycin assay

At 6HIV, embryonic cultures (generated as described above) were incubated with Schneider’s medium containing 25 μg/μl puromycin with or without 300 μg/μl cychlohexamide (CHX) for 30 min before fixation (Deliu et al., 2017). After fixation (same as described above) and 3 PBT washes, cells were stained for Puromycin, α-tubulin and HRP (see immunocytochemistry section).

### Imaging and data analysis

Data collection occurred over a period of 17 years and was made possible through a standardised data management scheme enabling us to retrieve and re-analyse well-documented original image files (Prokop, 2021b). Neurons were visualised using a compound fluorescence microscope (BX50WI or BX51; Olympus) and images of single neurons were captured using nijiBlueBox and the MatrixVision mvBlueFox3-M2 2124G camera at 100x magnification. Images were analysed using the FIJI/ImageJ 1.54p software: to determine the degree of MT disorganisation in axons we used the “MT disorganisation index” (MDI; Qu et al., 2017): the axon length (from cell body to tip of the most distant microtubule) was measured using the segmented line tool; area of disorganisation was measured using the freehand selection tool; this value was then divided by the the product of axon length multiplied with 0.5 μm (arbitrary axon diameter, thus approximating the expected area of the axon if it were not disorganised). All data were normalised against their respective controls. In each experiment, usually three slides per genotype were analysed aiming to image ∼30 isolated neurons per slide. Experiments were repeated at least once, data pooled. MDI data were usually not normally distributed but nevertheless plotted as mean ± SEM to avoid misleading median values of zero. Most experiments had only two groups and were assessed using Mann–Whitney Rank Sum tests, experiments with more then two group using Kruskal–Wallis one-way ANOVA with post hoc Dunn’s test. Means of single slides were used to generate super-plots (Lord et al., 2020) and assessed using standard t-tests. To assess ratios of tubulin/acetylated tubulin staining intensity ratios, 2-3 segments of MTs were drawn using the segmented line tool first in bundled regions then in curled regions of an individual neuron and the intensities of tubulin and acetylated tubulin (mean, min, max) were measured for each individual segment using the ROI window; the mean intensities of bundled segments and of curled segments in each neuron were averaged separately for each channel and their acetylated tubulin / total tubulin ratio determined. All statistical analyses were performed with Graphpad 10.2.2. Raw data files are available on figshare.com (https://doi.org/10.6084/m9.figshare.33112271). Figures were generated with Photoshop CS6, Illustrator CS and Canva Affinity.

### smFISH analyses

For smFISH, cells were fixed 20 minutes at RT with the same fixative and then permeabilised in 70% EtOH for 2 hours up to 2 days at 4°C. But we analysed the abundance of tubulin mRNAs and tubulin proteins. For tubulin mRNA we used single molecule fluorescent *in situ* hybridisation (smFISH; Orjalo et al., 2011) with specifically designed probes for *α1-* and *β1-tub* mRNA; mRNAs were counted along axons where single dots were easy to identify (in contrast to their clustered appearance in cell bodies). In control experiments, depending on culturing strategy and analysis method (see Suppl. Mat.), we counted mean mRNA numbers of 5 to 23 per axon and per tubulin isotype; both *α1*- and *β1-tub* smFISH probes proved specific because they showed a stark reduction in number when assessed in neurons deficient for the respective tubulin gene (Pinho-Correia, 2023).

### Reverse transcription and qRT-PCR analysis

To isolate RNA, 30 to 50 third instar larval brains were dissected in PBS and transferred into a tube containing 100 μl Trizol. Larval brains were homogenised with a pestle under the fume hood, 100 μl of chloroform were added and closed tubes were vortexed for a minute, left to incubate at room temperature for 3-5 min and then centrifuged (4°C, 12,000 g, 20 min). 100μl of the upper, clear, RNA-and DNA-containing phase was added to 100 μl pure isopropanol and centrifuged (4°C, 12,000 g, 10 min). The supernatant was discarded and the pellet washed twice with 70% ethanol, adding 400 μl for the first washing step and 250 μl for the second, each step followed by centrifugation (4°C, 12,000 g, 5 min) and discarding the supernatant. The pellet was dried and resuspended in 50-60 μl double-distilled, RNA-free water (ddH_2_O).

Reverse transcription was carried out using the QuantiTect reverse transcription kit (Qiagen). 500 ng RNA (measured using a NanoDrop by Thermo Scientific according to the manufacturer’s instructions) and 1 μl gDNA wipeout buffer (provided as kit component) were added to 7μl ddH_2_O in 0.2 ml PCR 8-Strip Tubes (Starlab) and incubated for 2 min at 42°C. 3μl of a mastermix, containing 2 μl Reverse-Transcription-Buffer, 0.5 μl Primer-Mix and 0.5 μl Reverse-Transcriptase were added and the sample was incubated in a thermocycler (Thermal Cycler C1000 by BioRad) for 15 min at 42°C. The Reverse-Transcriptase was then inactivated at 95°C for 3 min. The sample was filled up with 90μl of ddH_2_O and stored at -20°C.

cDNA samples were analysed using qPCR for which two 20 μl master mixtures were prepared: the first contained 10 μl of SYBRGreen Supermix, including fluorescent dye and 1 μl of 5 μM forward and reverse primer mix (lyophilised primer diluted according to the certificate of synthesis) for the gene in question (see table below). The second was composed of 1 μl cDNA from the stored sample described above and 8 μl ddH2O. The master mixes were prepared in 0.2 ml PCR 8-Strip Tubes and then pipetted into 96 well plates in doublets. The qPCR reaction was carried out using the BioRad® IQ5 Multicolor Real-time PCR detection system. As reference gene we used primers for the RNA of *rp49* encoding Ribosomal protein L32 (Hahn et al., 2013).

The following temperature and cycle protocol was used:

| Cycle step | Temperature | Time | Cycles |
| --- | --- | --- | --- |
| Initial denaturation | 95 °C | 2 min | 1 |
| Denaturation | 95 °C | 15 s |  |
| Annealing | 60 °C | 15 s | 45 |
| Extension | 72 °C | 1 min |  |
|  | 95 °C | 15 s |  |
| Melt curve | 65 °C - 95 °C<br>(0.3 °C increment) | 15 s | 1 |

The following primers were designed using Perlprimer software (https://perlprimer.sourceforge.net):

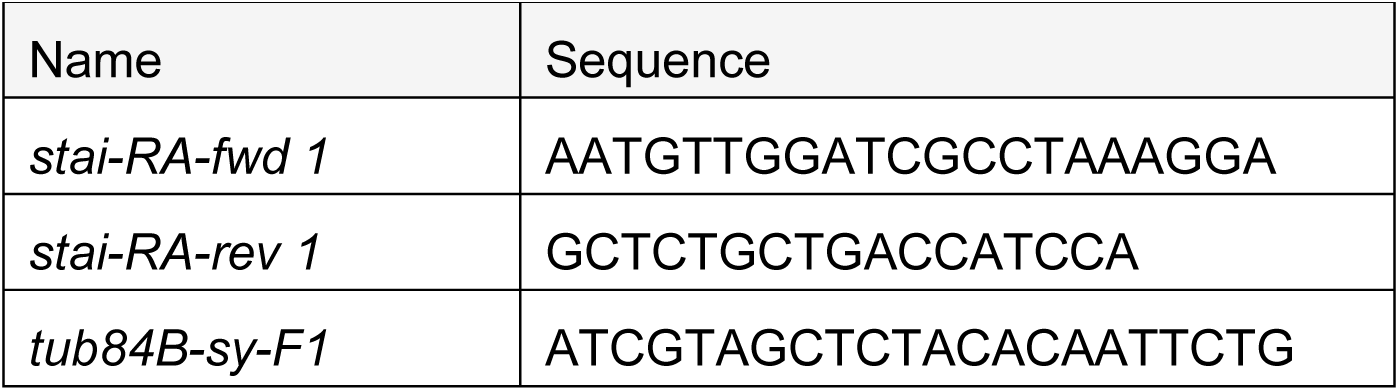
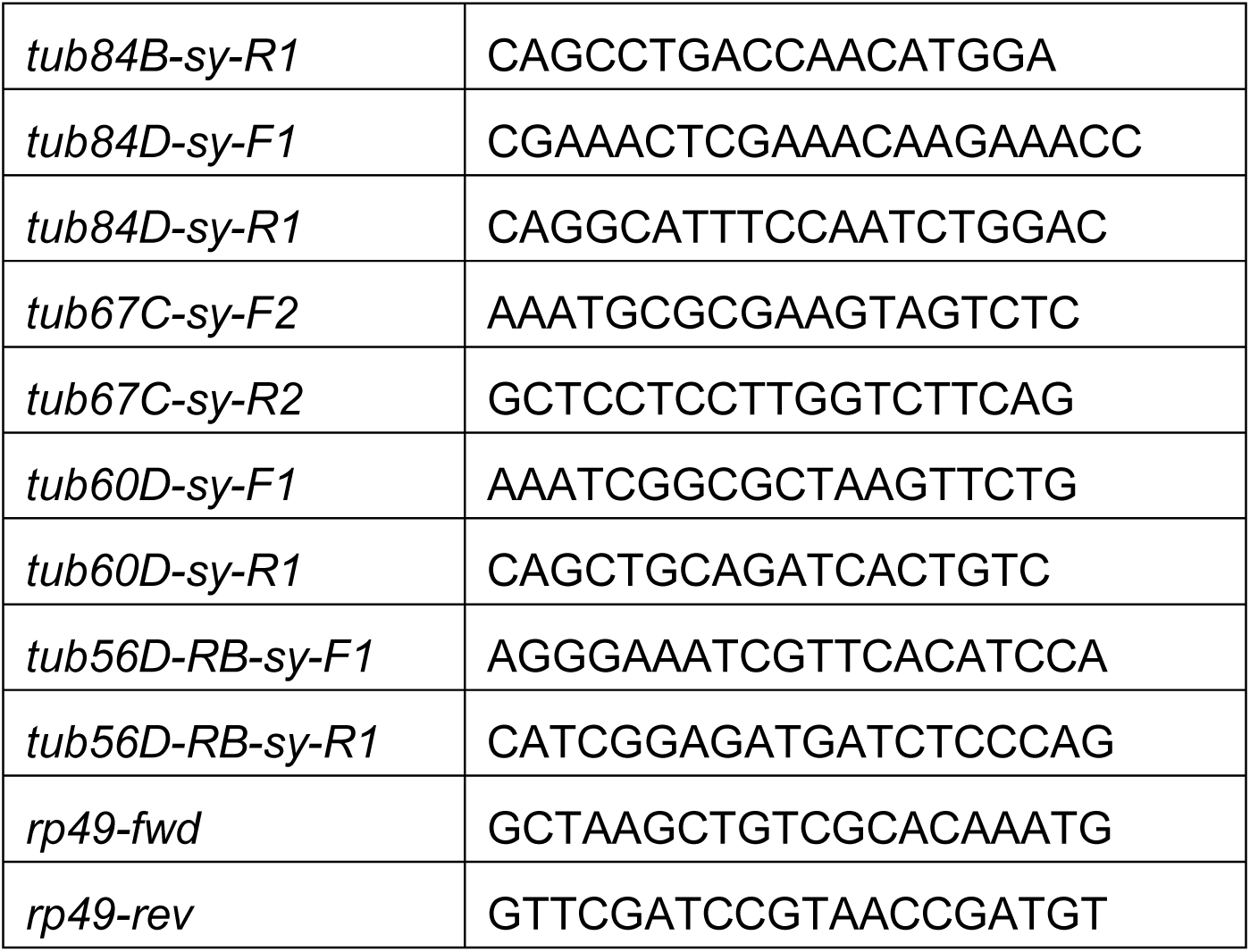

### Live imaging procedures

For long term imaging, embryos were collected, dechorionated and stage 11 embryos collected by auto-fluorescence pattern. 36 embryos were prepared for primary neuronal cultures (Voelzmann and Sánchez-Soriano, 2022) and cells re-suspended in 200 µl supplemented growth medium. The cell suspension was transferred to a 35mm ConA-coated glass bottom MatTek dish and left to adhere to the dish for 30 minutes. The dish was gently shaken to dislodge non-adhering cells, aspirated and then gently rinsed with 400 µl pre-warmed medium. The medium was then replaced with 3 ml pre-warmed growth medium (to buffer temperature fluctuations, mitigate effects of evaporation) with or without 100 nM SiR-Tubulin. The dish was sealed with Vaseline and a 40 mm coverslip to reduce evaporation of the medium during imaging (Voelzmann and Sánchez-Soriano, 2022). Samples in imaging chambers were transferred to pre-warmed imaging chambers and left to equilibrate for 15 minutes. 2 hours after plating, neurons were imaged.

The long-term live imaging was executed either on a 3I Marianas Yokogawa CSU-X1 spinning disc confocal system with infinite focus stabiliser or a Zeiss LSM880 (Plan Apochromat 63x/1.4 Oil DIC M27 lens) with Definite Focus 2 image stabilisation and environmental chambers. Environmental chambers were pre-warmed and equilibrated to 26°C for 45 minutes before samples were added. Samples were left to equilibrate for 15 minutes. Due to the unpredictability of the formation of areas of unbundling and direction of axonal growth, imaging required a large field of view (4×4 tiles or multi-position) while at the same time maintaining sufficient magnification and resolution to identify the formation of areas of microtubule unbundling. Samples were imaged as multi-position 4D stacks for 20 to 22 hours.

The applied imaging setup when using the 3I Marianas Yokogawa microscope: Jupiter-mCherry labelled cells were imaged for ∼15 hours as 4D stacks (3 z-planes, 0.34 µm steps) at multiple positions on the MatTek dish with a timelapse interval of 2 minutes, an exposure time of 500ms, 60% laser power (561 nm) and again of 2. On the Zeiss LSM880 microscope, Jupiter-mCherry labelled samples were imaged as image stacks (5 z-planes, 0.35 µm steps) of 16 tiles (4×4) with 3% laser power (561 nm), emission detection at 578-696nm, and 4x line average every 5 minutes. SiR-Tubulin labelled samples were imaged as image stacks (5 z-planes, 0.6 µm steps) of 16 tiles (4×4) with 2% laser power (633 nm), emission detection at 615-695 nm, 2x line average every 5 minutes. Even with definite focus, we noticed a z-shift of ∼6 µm over the 19-22 hours imaging period and compensated by adapting the z-stack coordinates.

### Analysis of live movies

*Analysis of live movies*: 4D stacks were converted to maximum intensity projection time courses. Criteria that were applied for cell analyses were: 1) neurons grew axons at least twice as long as the soma diameter; 2) neurons survived the imaging time course; 3) The entirety of axons stayed in focus and were visible throughout the time course of imaging.

*Measuring the size of areas of disorganised MT curling over time*: areas of MT unbundling were identified at timepoints when they were most obvious and then traced back to the timepoint of their first initiation which was defined as timepoint 0 for this disorganised region. The area of disorganisation (µm^2^) was determined in intervals of 30 min and averaged for multiple regions (n = 7 to 26). Due to the variability of the timepoint of curl initiation, the time course gets more data-sparse towards later timepoints.

*Onset of disorganisation*: to determine the onset of disorganisation, the timepoint of the first appearance of a region of microtubule unbundling was determined relative to the imaging start (∼45 minutes after plating primary cultures).

*Origin of disorganisation*: To determine the origin of disorganisation, regions of disorganisation were traced back to the first unbundling. Unbundling starting at the soma and spreading from there were classed as ‘soma’, unbundling starting at the tip of the growing axon at the time of first appearance were classed as ‘tip’, unbundling starting in the axon shaft classed as ‘shaft’ and unbundling originating during the formation of a branch were classed as ‘branch’. The number of events for each classed was scored.

Number of disorganised regions: The number of unbundled regions along the axon (irrespective of their size) was counted throughout the imaging-lifetime. Only regions on the main extension but not along branches were counted.

---

**Fig. S1.**
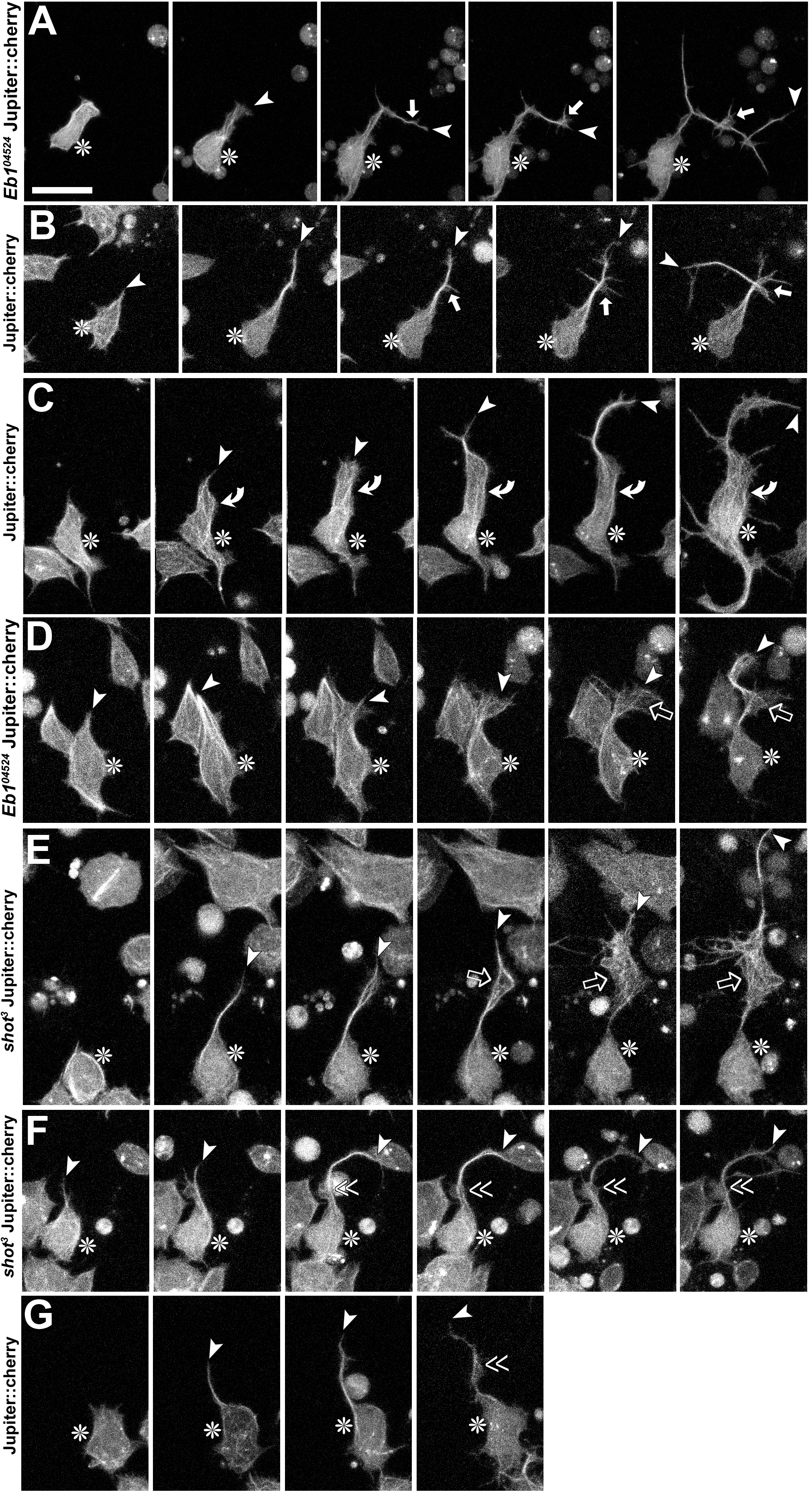
Sites of MT-curl initiation as observed during live imaging. Each row represents selected images from a respective film sequence of a neuron with a given genotype and labelled as indicated on the left (see also Fig.3). Asterisks indicate cell bodies, arrow heads axon tips, white arrows curl initiation at branch points, curved arrows initiation close to the soma, open arrows in growth cones, and double chevrons along the axon shaft.

**Fig. S2.**
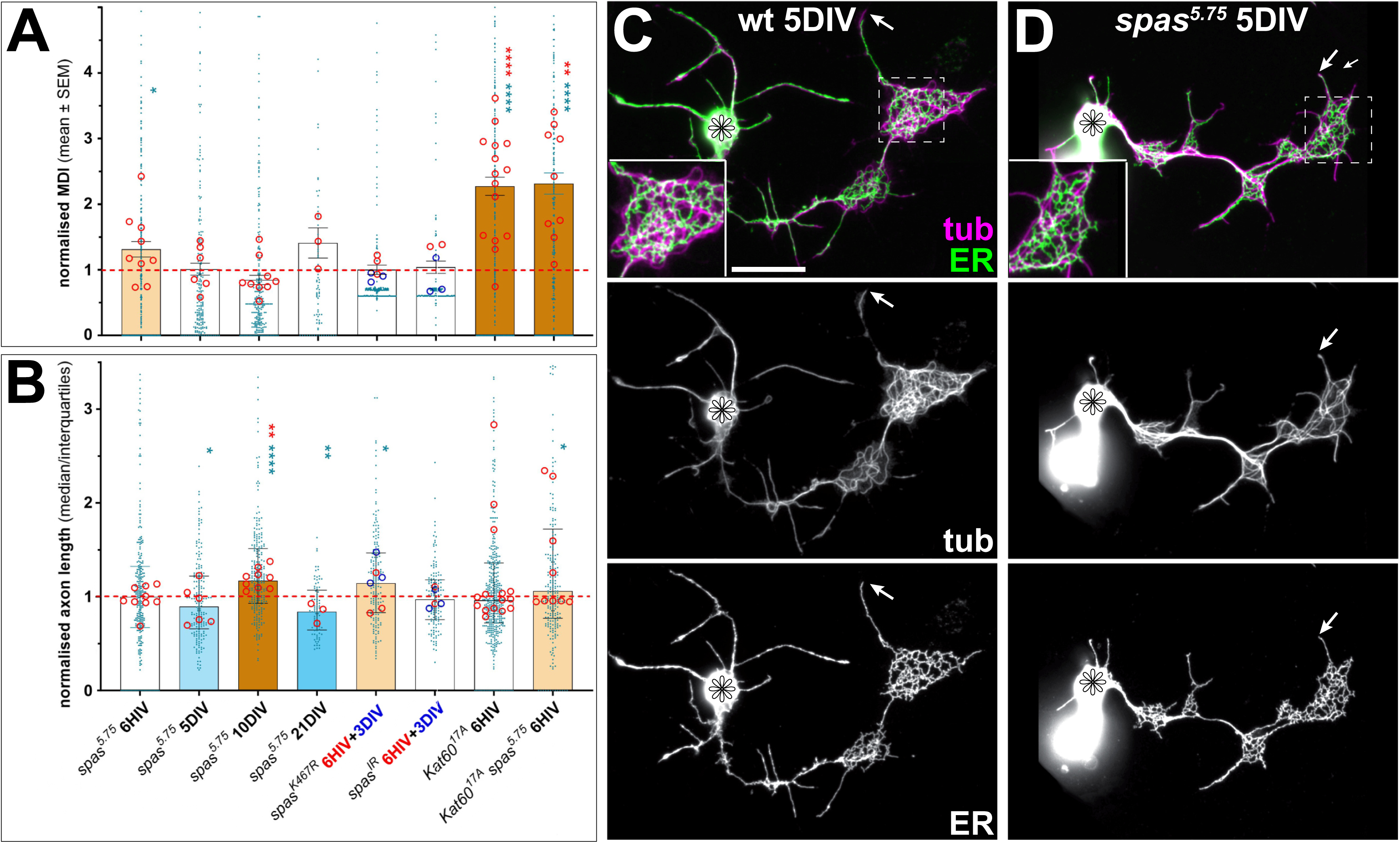
Analyses of MT severing proteins reveals that Katanin is a suppressor of MT curling. **A,B)** Analyses of MT-curling (MDI; A) and axon length (B) in different mutants, presented with the same logic as explained in Figs. 4 and 6 (for detailed data see Tab.1). Analysis of neurons after 6 hours *in vitro* (HIV) or after 3, 5, 10 or 21 days *in vitro* (DIV) which are homozygous for the *spas^5.75^*null allele (confirmed by RT-PCR; data not shown), express the *spas^K467R^*point mutation (Orso et al., 2005) in wild-type background, display spas^IR^-mediated knock-down, carry the *Kat60^17A^* null allele or a combination of *spas^5.75^* and *Kat60^17A^* in homozygosis. Graphs are organised as explained in Fig.6; for *spas^K467R^* and *spas^IR^* data from 6HIV (red) and 3DIV (blue) experiments are combined in one graph, with circles indicating repeats colour coded by experiment. **C,D)** A wild-type (C) and *spas^5.75^* mutant neuron at 5DIV stained for tubulin and the ER marker Rtnl1::YFP, shown in combination in colour or as single channels in greyscale (as indicated); both neurons show an area of MT curling (inset twofold magnified) where the ER displays the same characteristic 3-way junctions known from non-neuronal cells; this would suggest that expected ER-regulating roles of Spastin (Öztürk et al., 2020) are at most subtle. Scale bar in C represents 20µm in C and D. Raw data files are available on figshare.com (https://doi.org/10.6084/m9.figshare.33112271; see in Tab.S1 folder).

**Fig. S3.**
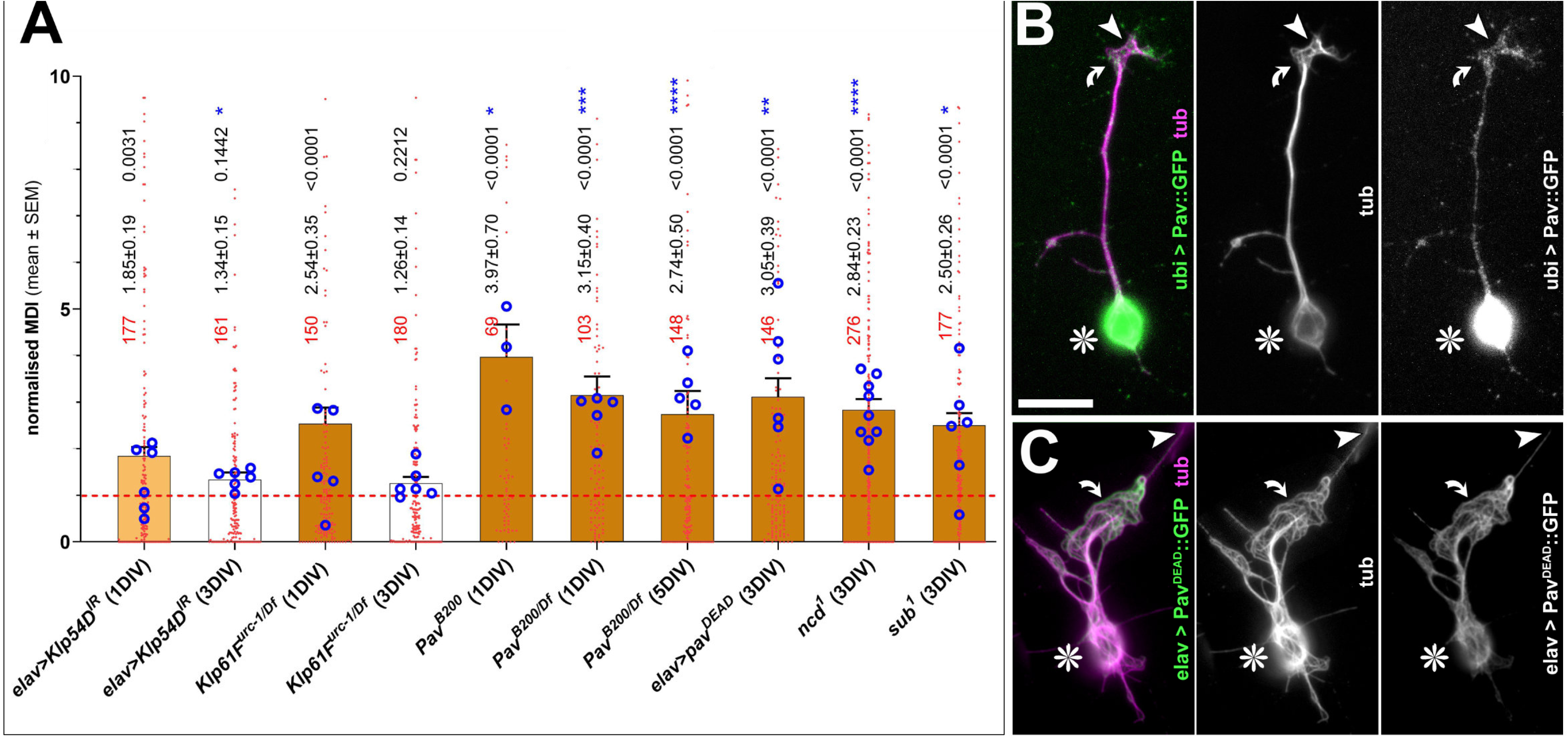
Analysis of mitotic kinesins. **A)** Analyses of MT-curling (MDI) in different mutants, presented with the same logic as explained in Figs. 4 and 6 (for detailed data see Tab.1). Neurons display *elav-Gal4*-driven knock-down of Klp54D, carry the *Klp61F^urc-1^* null allele over a deficiency (*Df*) uncovering the *Klp61F* locus, carry the *pav^B200^* null allele either in homozygosis or over deficiency, or display *elav- Gal4*-driven expression of the Pav^DEAD^ rigour-type variant (Minestrini et al., 2002). Cells were cultured for 1, 3 or 5 days *in vitro* (DIV) as indicated. **B,C)** Neurons stained for tubulin (tub, magenta) and GFP (green), with the neuron in B displaying Pav::GFP expression driven by a ubiquitin promotor not well aligning with MTs, and neurons in C showing an *elav-Gal4*-driven rigour mutant version of Pav (Pav^DEAD^::GFP) which tightly decorates axonal MTs as is typical of rigour mutants (Blangy et al., 1998; Klumpp et al., 2003; Minestrini et al., 2002; Nakata and Hirokawa, 1995). Asterisks indicate cell bodies, arrow heads axon tips, curved arrows areas of MT curling, and the scale bar in C represents 20µm in C and D. Raw data files are available on figshare.com (https://doi.org/10.6084/m9.figshare.33112271; see in Tab.S1 folder).

**Fig. S4.**
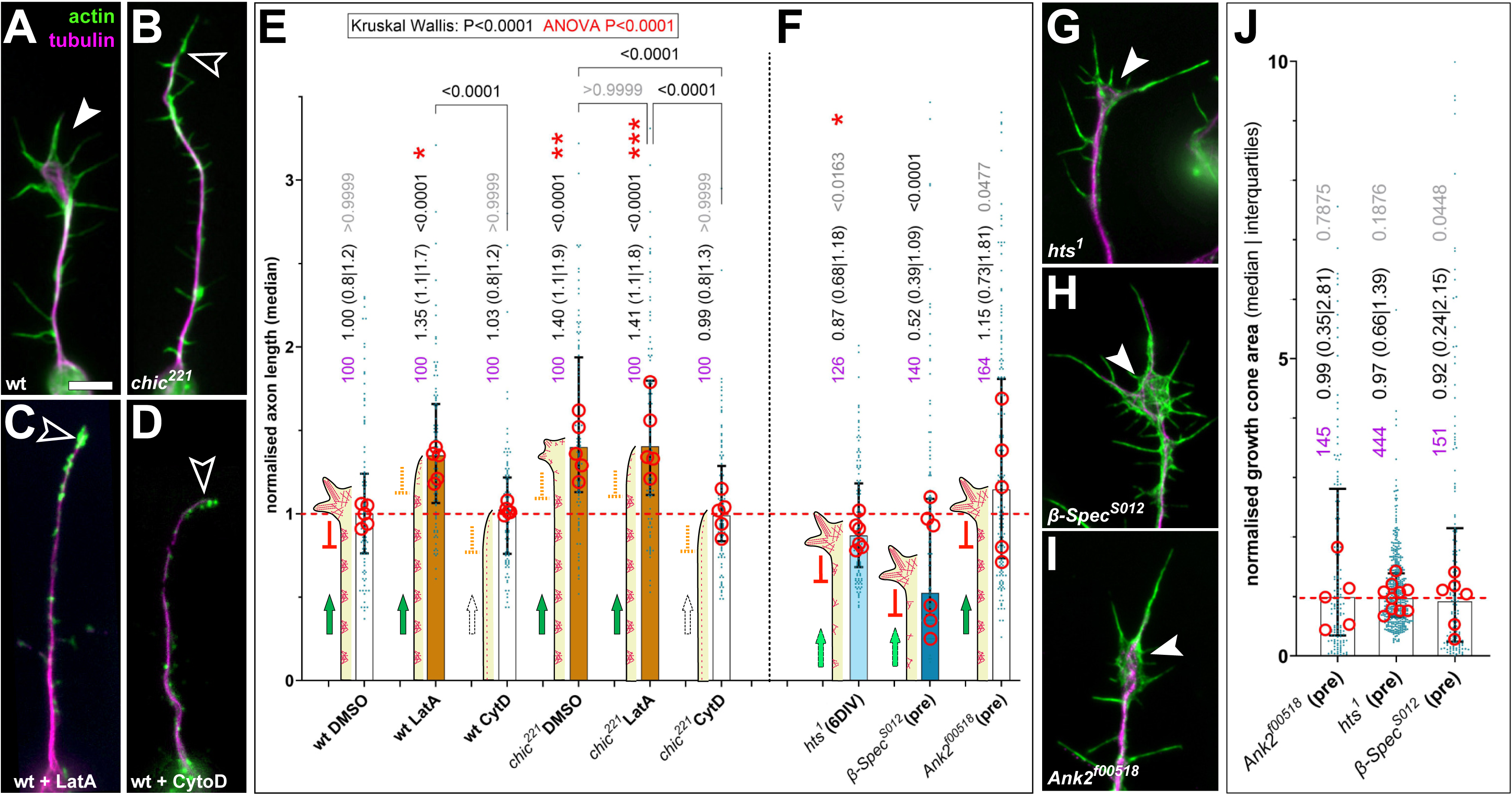
Different roles for F-actin networks in axon shafts versus growth cones. **A-C, G-I**) Images of axons with (white arrow heads) or without (open arrow heads) prominent growth cones at their tips; all neurons are labelled for tubulin (magenta) and actin (phalloidin stain in green) and, as indicated bottom left in each image, are of different genotypes or treated with the F-actin inhibitors latrunculin A (LatA) or cytochalasin D (CytD), which both reduce growth cone size but only CytD eliminates F-actin rings in the axonal shaft (Qu et al., 2017); the scale bar in A represent 10 µm in A-D and 7.5 µm in G-I. **E**) One experiment with 6 groups including wild-type (wt) and *chic^221^* homozygous mutant neurons at 6 hours in vitro, treated with vehicle (DMSO), LatA or CytD, as indicated; sample numbers of analysed neurons shown in blue, described via median and interquartile ranges and assessed by Kruskal-Wallis (box at the top) and Dunn’s multiple comparisons test (p values given); experimental replicates from two repeats are indicated by red circles and assessed by ANOVA (in box at the top) and Dunnett’s multiple comparisons indicated by red asterisks (* P ≤ 0.05; ** P ≤ 0.01; *** P ≤ 0.001; **** P ≤ 0.0001). The schematics on the left of each bar illustrate our proposed interpretation where the presence of large growth cones slows down axon growth (red T-bar) and the presence of actin rings (magenta mesh structures) promotes axon growth (green arrow); loss of different F-actin networks reduces these regulatory impacts (stippled T-bars or arrows) shifting the net-growth outcome. **F**) A similar plot as in E for loss of other actin regulators, none of which affects growth cone size (shown in G-I and plotted in **J**), of which loss of Hts and β-Spectrin reduce F-actin rings in the axon shaft, but not Ankyrin 2 (Qu et al., 2017). Raw data files are available on figshare.com (https://doi.org/10.6084/m9.figshare.33112271).

**Fig. S5.**
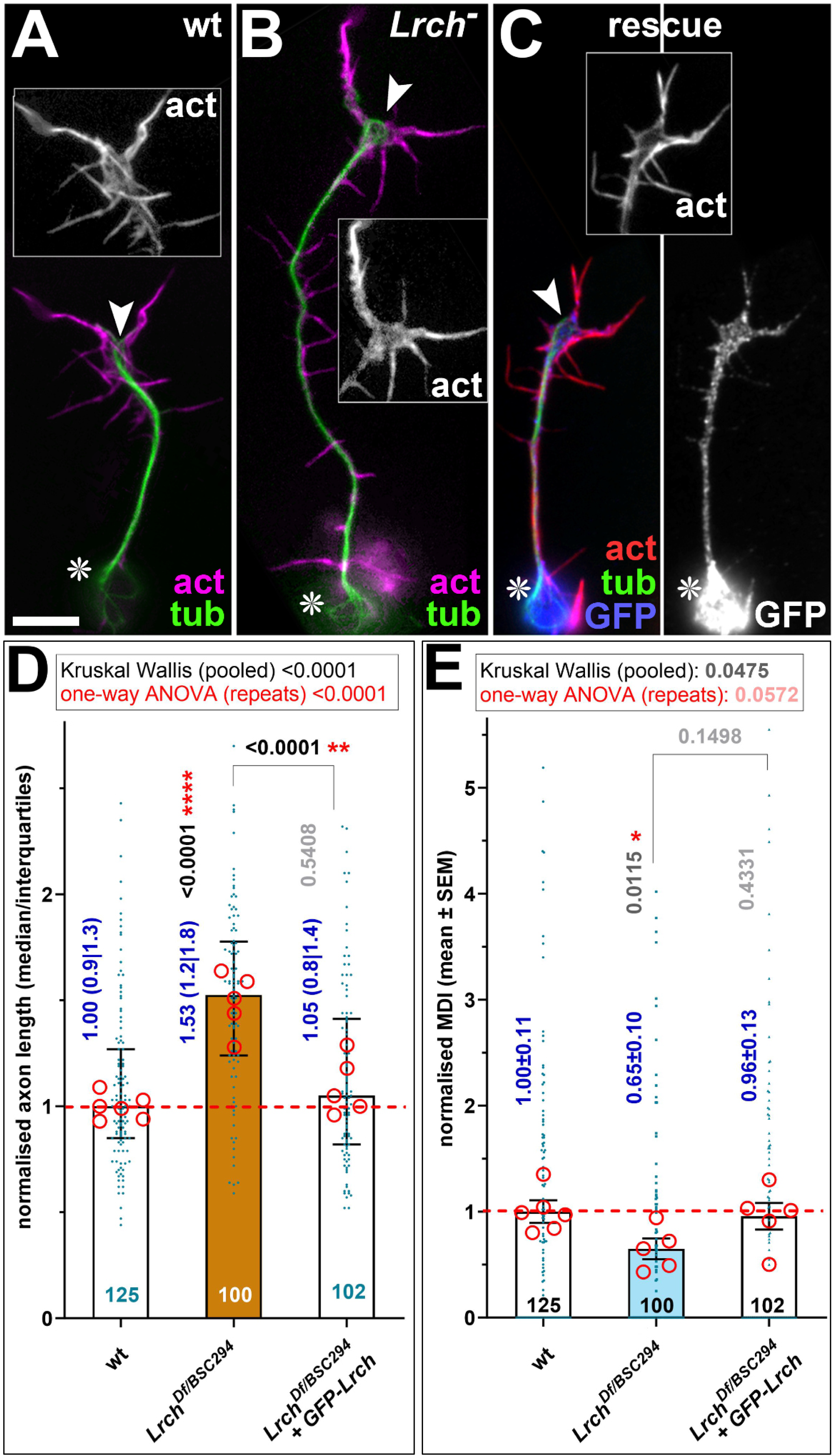
Lrch as a new axon length regulator. **A-C)** Neurons at 6 hours *in vitro* (HIV) stained for actin (act), tubulin (tub) and GFP, as indicated in colour-code bottom right; neurons are A) wild-type (wt), B) carry the small deletion *Lrch^Df^* over another larger deletion uncovering the locus (*BSC294*; same results were obtained using *Lrch^Df^*in homozygosis; Beaven, 2012), or C) display the same *Lrch^Df/BSC294^* mutant background but also express GFP::Lrch driven by *sca-Gal4* (rescue); insets show enlarged greyscale views of actin staining in growth cones, and the figure on the right in C) a single-channel greyscale view of GFP::Lrch expression; asterisks indicate cell bodies, arrow heads the tips of axons, and the scale bar in A represents 15µm in all images except insets. **D,E)** Quantification of data obtained from neuronal cultures shown in A-C comprise analyses of the normalised MDI (A) and normalised axon lengths (B) presented with the same logic as explained in Fig. 6; since only data for *Lrch^Df/BSC294^*are shown in Tab.S1, detailed information is provided in the graph: numbers at bottom of bars show how many neurons were analysed, blue numbers indicate median with interquartile ranges in D) and mean ± SEM in E); grey/black numbers show significance values of pooled data established via Mann-Whitney, red asterisks the results from t-test analyses of replicates (red circles; * P ≤ 0.05; ** P ≤ 0.01; *** P ≤ 0.001; **** P ≤ 0.0001) and the box at the top shows analyses across the three groups for pooled data (Kruskal Wallis) and replicates (one-way ANOVA). Raw data files are available on figshare.com (https://doi.org/10.6084/m9.figshare.33112271).

**Fig. S6.**
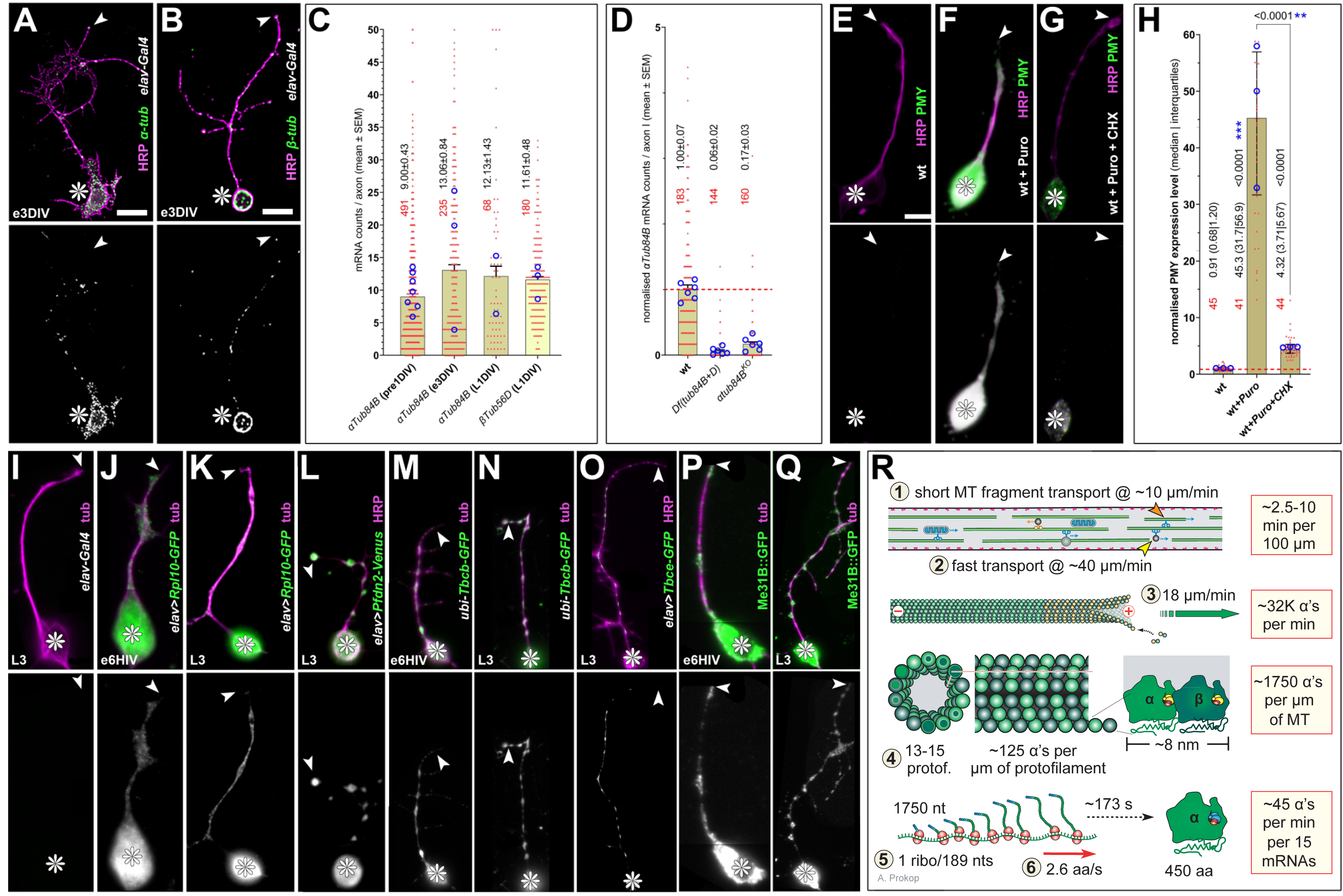
Local translation of tubulins appears to occur in axons but is likely negligible. Photographic images show that examples of various factors required for local tubulin biogenesis are present in axons; they show neurons derived from embryos (indicated by suffix ‘e’ bottom left of images) cultured for 3 days in vitro (DIV) or 6 hours in vitro (HIV), or derived from late larval CNSs (L3) and cultured for 1 day. As indicated on the right of each image, neuronal morphology is shown in magenta stained with the surface marker HRP or with anti-tubulin (tub), whereas specific sub-components are shown in green: **A**) α-tub, smFISH analysis of *αTub84B* (α1-tubulin) mRNA; **B**) β-tub, smFISH analysis of *βTub56D* (β1-tubulin) mRNA; **E-G**) PMY, anti-puromycin staining indicating puromycin incorporation into proteins translated during the incubation period in the absence of presence of puromycin (puro) in the culture medium; puromycin resembles the 3′ end structure of amino-acylated tRNA (aa-tRNA) with a modified adenosine base covalently linked to tyrosine (Tercero et al., 1996; Yarmolinsky and Haba, 1959) so that its incorporation into the C-terminus of elongating nascent peptide chains irreversibly and prematurely terminating translation and release of the peptide from ribosomes into the cytoplasm (Semenkov et al., 1992; Starck and Roberts, 2002); cycloheximide (CHX) inhibits active translation including puromycin insertion (Deliu et al., 2017) and serves therefore as a control. **I**) a typical control neuron without targeted expression; **J**) Rpl10-GFP, fluorescently tagged ribosome component; **K**) Pfdn2-Venus, fluorescently tagged prefoldin chaperone complex component (first steps of tubulin maturation; Pinho-Correia and Prokop, 2023); **M-O**) Tbcb-GFP/Tbce-GFP, fluorescently tagged tubulin-binding cofactors B and E (3^rd^ step of tubulin maturation); **P**,**Q**) Me31B, mRNA-binding protein typical of P-bodies; ‘*elav>*…’ indicates that UAS-constructs (in green) were driven by *elav-Gal4*, ‘ubi-…’ indicates a fusion construct with ubiquitin promoter. The scale bar in A represents 10µm in A,C,J, 15µm in B,E,F and 20µm in G-I,K. **C**) Quantitative analysis plotting the number of smFISH-labelled dots detecting α1-(darker columns) or β1-tubulin mRNAs (lighter column) in axons of neurons pre-cultured from embryos (pre), cultures for 3 days from embryos (e3DIV) or from larval CNSs for 1 day (L1DIV); single data are shown as red dots, sample numbers in red and the mean ± SEM is indicated; mean values of replicates are shown as blue circles. **D**) Validation experiment with neurons pre-cultured from wild-type (wt) or mutant embryos deficient for *αTub84B* demonstrating the specificity of the smFISH probe used in our experiments. **H**) Quantification of puromycin intensity measured with NeuronJ in the absence or presence of puromycin (Puro) and validated with cycloheximide (CHX) application; data for single neurons are shown as red dots, sample numbers in red and described as median and interquartile ranges; medians of replicates are shown as blue circles; Kruskal-Wallis analysis reveals P<0.0001 with Dunn’s multiple comparisons test results shown in black and t-test results of replicate medians as blue asterisks (* P ≤ 0.05; ** P ≤ 0.01; *** P ≤ 0.001; **** P ≤ 0.0001). **R**) Calculations based on data from the literature as indicated by encircled numbers: (1) (Dent et al., 1999), (2) (Twelvetrees, 2020), (3) (Zwetsloot et al., 2018), (4) (Prokop, 2020), (5) (Hendrickson et al., 2009), (6) (Riba et al., 2019), with conclusions shown emboxed on the right; from bottom to top: ∼15 mRNAs (see C) produce only ∼45 proteins per minute considering proposed data for ribosomal propagation and density on the 1750 nucleotide-long *α1-tubulin* mRNA; a 1 µm-long stretch of MT contains ∼1750 α1-tubulins considering on average 14 protofilaments per MT and 8 nm length of α/β-heterodimers; the average polymerisation speed of MTs would require ∼32,000 α-tubulins per minute; fast axonal transport of tubulin hetero-dimers or slower transport of MT fragments could reach the tip of typical axons in *Drosophila* cultures within 10 minutes (Baas, 2002; Baas and Buster, 2003). Especially transported fragments could be used as tubulin source through disassembly and recycling processes (Kortazar et al., 2007; Nogales and Wang, 2006; Nolasco et al., 2021). In contrast to flies, local tubulin biogenesis might be far more important in the much longer vertebrate axons. Raw data files are available on figshare.com (https://doi.org/10.6084/m9.figshare.33112271).

**Fig. S7.**
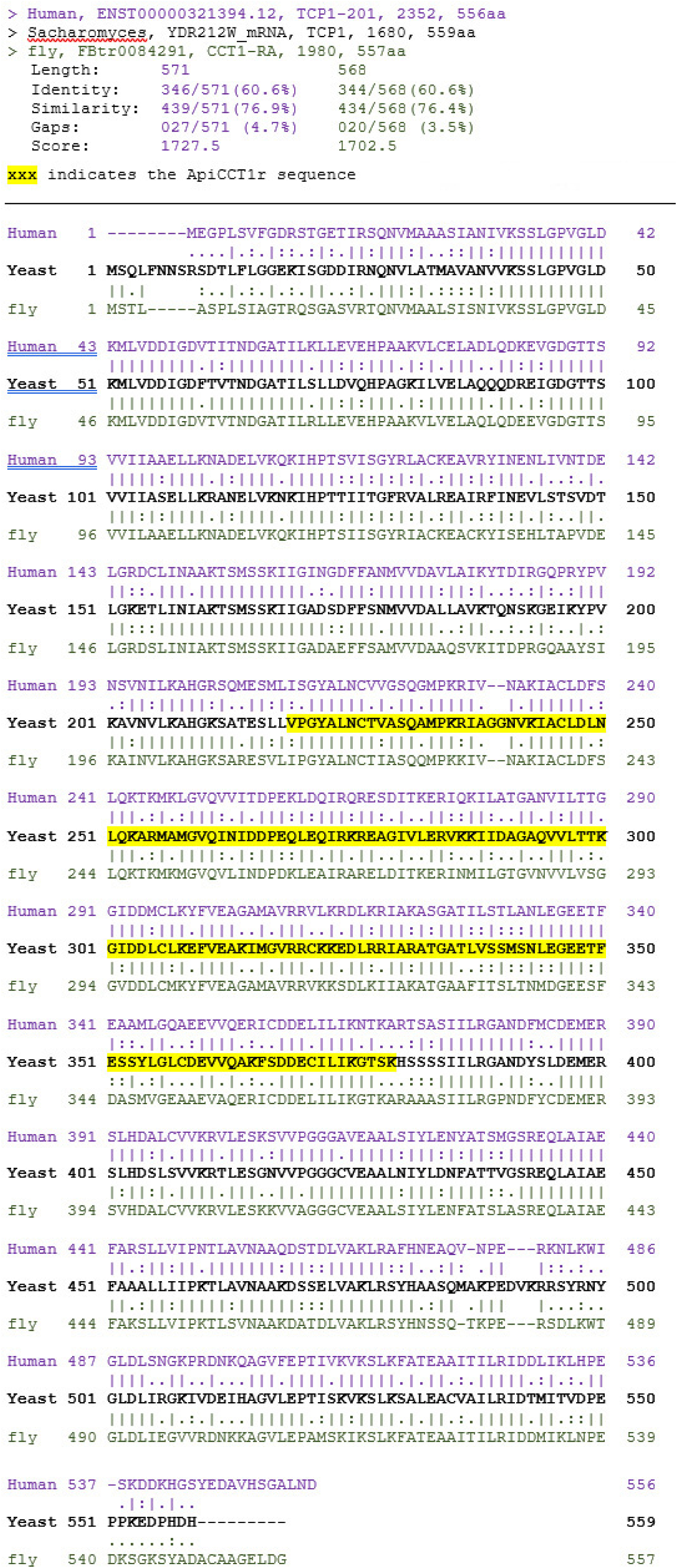
CCT1 protein sequences of yeast, fly and human show homology of the ApiCCT1r sequence known to ameliorate protein aggregates in Huntington’s disease models. Analyses were performed with https://www.ebi.ac.uk/jdispatcher/psa/emboss_needle using the indicated amino acid sequences of yeast (black) aligned with human (blue) and fly (green). The ApiCCT1r sequence (Sontag et al., 2013) is highlighted in yellow and appears well-conserved in human and fly (vertical lines indicate identity, dots and colons similarity; percentages for the overall sequence comparison are given at the top).

**Fig. S8.**
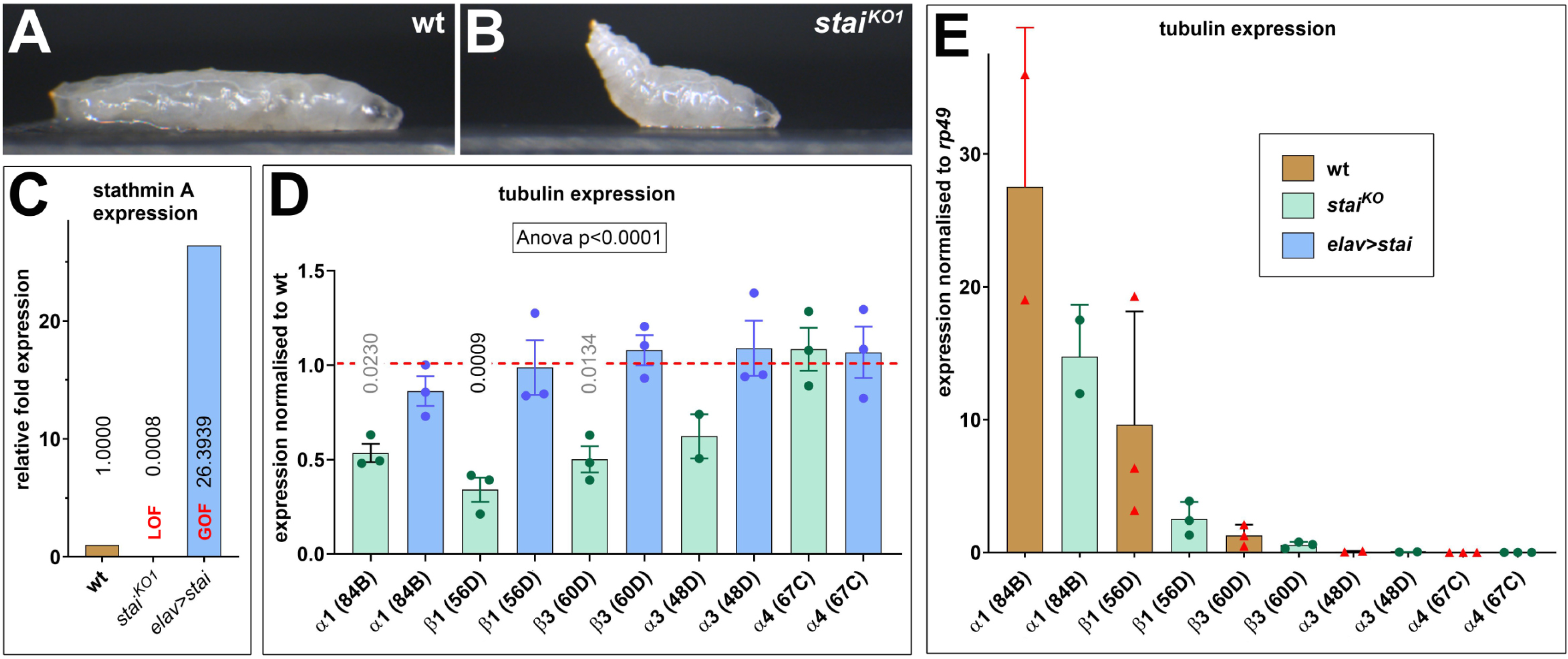
α1 and β1 tubulin are the only prominent isotypes in the fly CNS and their levels are maintained by Stathmin. **A,B**) Larvae homozygous for the *stai^KO1^* mutation show the same ‘tail-flip’ phenotype as reported previously for other mutant alleles of stai (Duncan et al., 2013). **C**) Quantitative RT-PCR analyses show the loss of *stai* transcripts in *stai^KO1^* mutant larval CNS (LOF, loss of function) and a stark increase upon *elav-Gal4*-driven over-expression of *stai* (GOF, gain of function). **D**) Quantitative RT-PCR analyses of late larval CNSs show that loss of Stai (grey bars) causes a reduction of most tubulin isotype mRNA; data are normalised to wild-type (stippled red line) and assessed by ANOVA, with Dunnett’s multiple comparisons results provided in the graph. **E**) Normalisation of data to mRNA of the housekeeping gene *rp49* (Ribosomal protein 32) shows that α1- and β1-tubulin are the dominating tubulin isotypes in the larval CNS, as is supported by micro array analyses (‘FlyAtlas2’: <u>motif.mvls.gla.ac.uk/FlyAtlas2</u>; Krause et al., 2021) and available RNAseq data (‘modEncode’: www.modencode.org/publications/about/index.shtml; Celniker et al., 2009). Raw data files are available on figshare.com (https://doi.org/10.6084/m9.figshare.33112271).

**Fig. S9.**
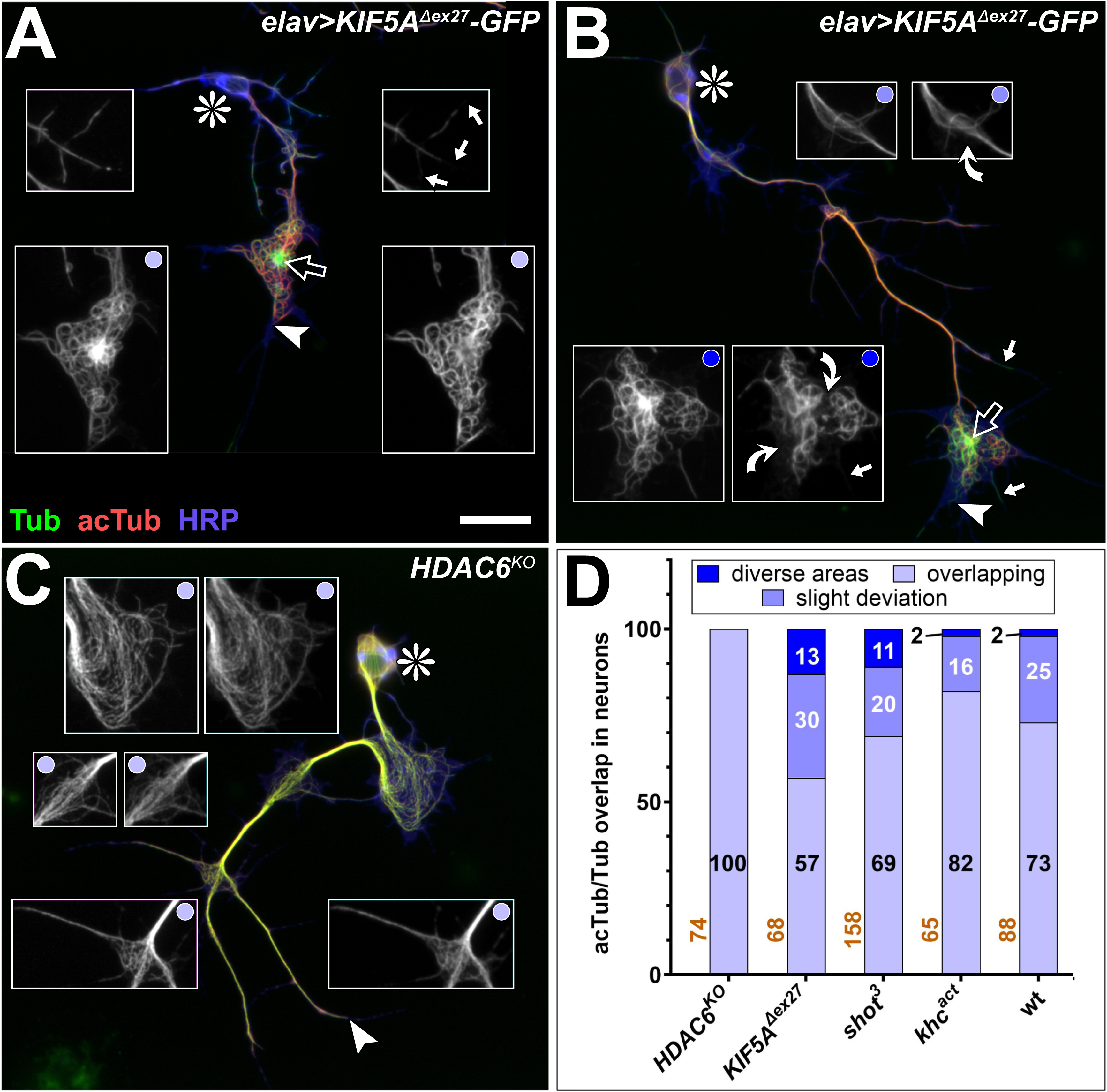
Impacts of genotype and MT-curling on MT acetylation. **A-C**) Primary neuron with areas of MT-curling which are of different genotypes (indicated top right) and stained for tubulin (green), acetylated tubulin (red) and the neuronal membrane marker HRP (blue). 1.5-fold magnified pairs of insets show regions of interest as single channel images of total tubulin (left) versus acetylated tubulin (right). Symbols indicate cell bodies (asterisks), axon tips (arrow heads), aggregates of KIF5A^Δex27^::GFP (open arrows); curved arrows show examples of ‘diverse areas’ in insets with dark blue dots or examples of ‘slight deviation’ in insets with middle-blue dots; insets with light blue dots show examples of ‘overlapping’ staining and white arrow heads point at MTs in filopodia lacking acetylation at their distal segments. Scale bar in A represents 20 µm in all images. **D**) Quantification of the fractions of neurons per genotype that display either of the three classifications of staining overlap or deviation that were illustrated in A-C.

**Tab. S1.**
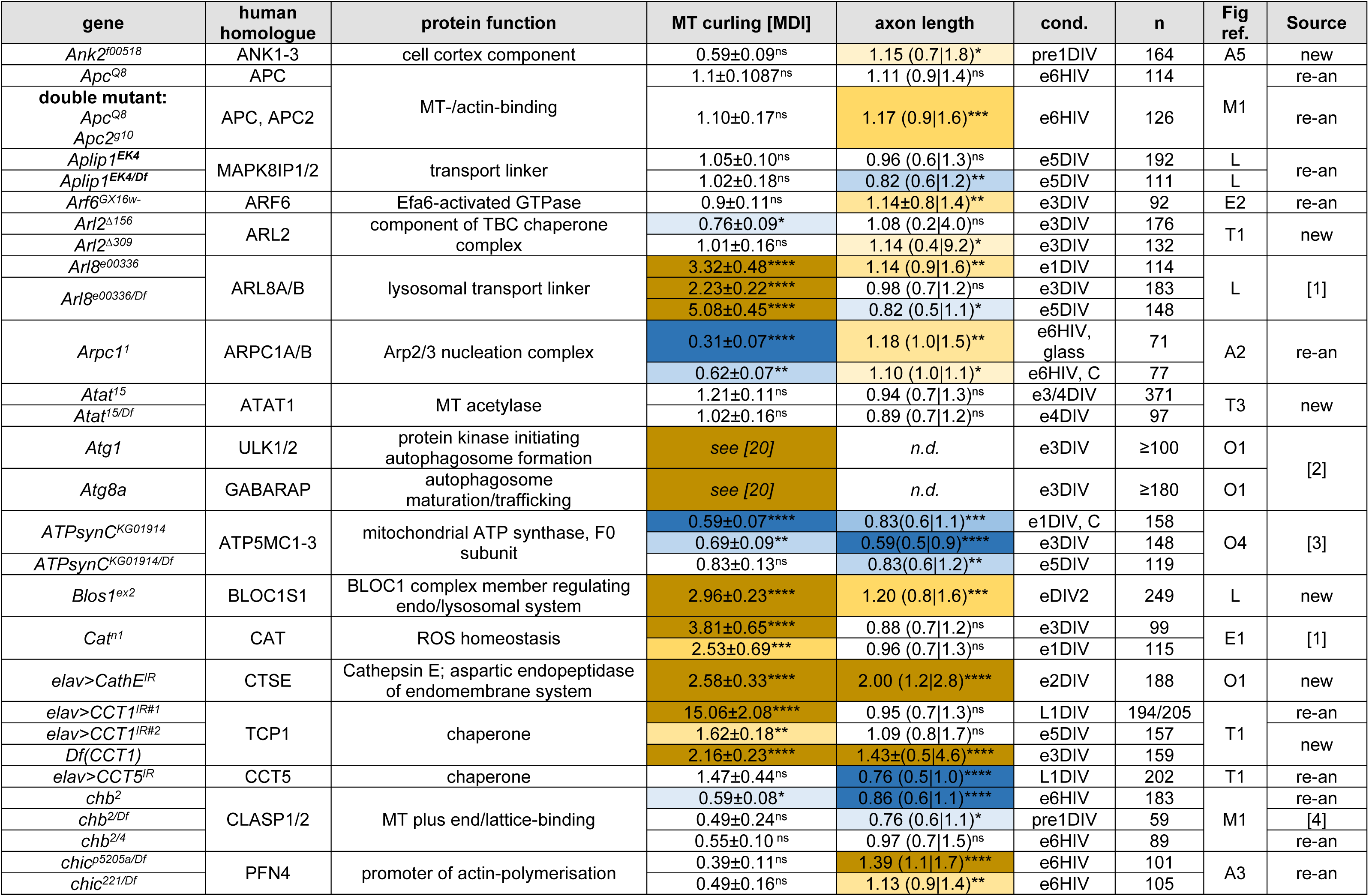
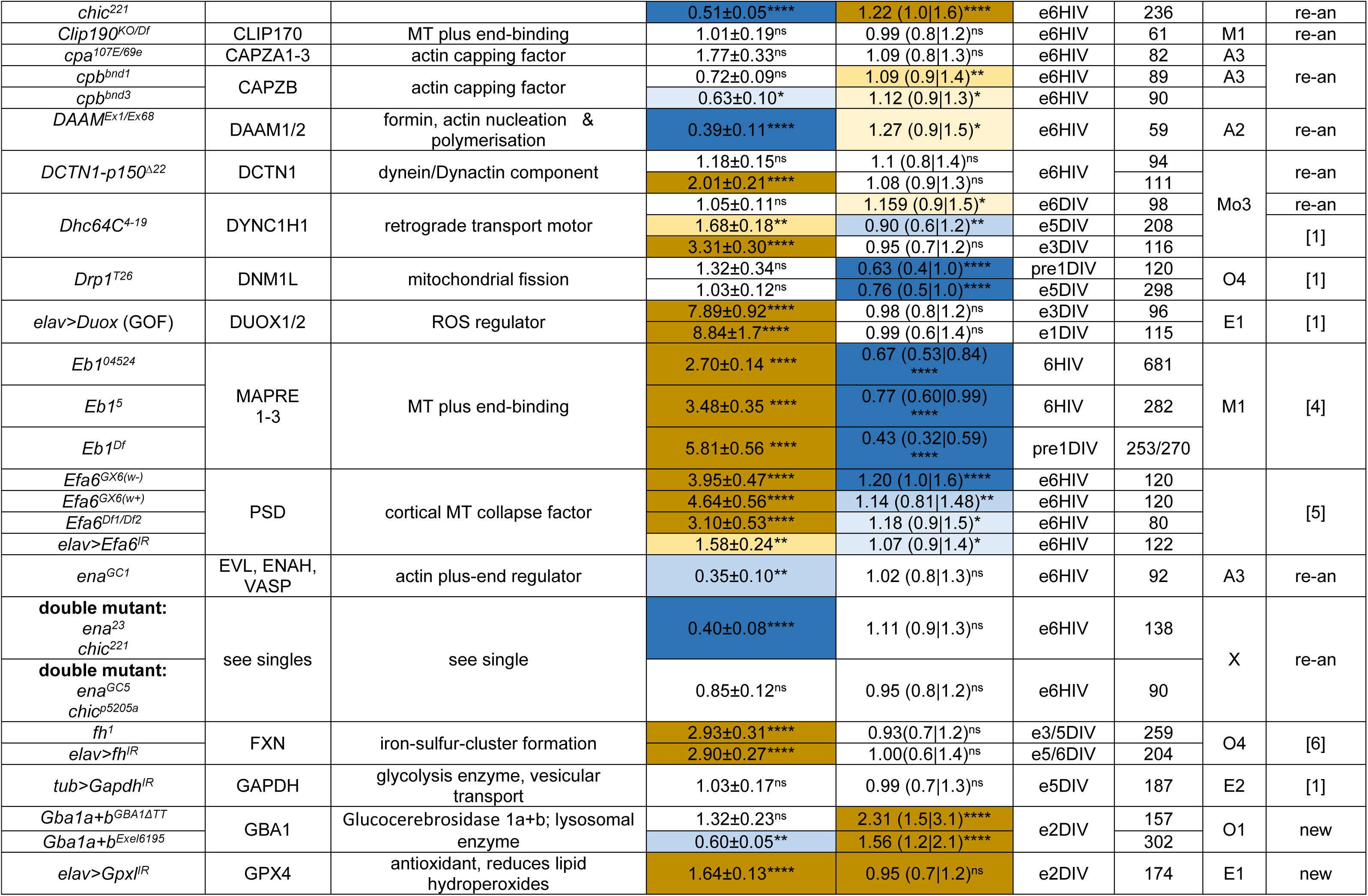
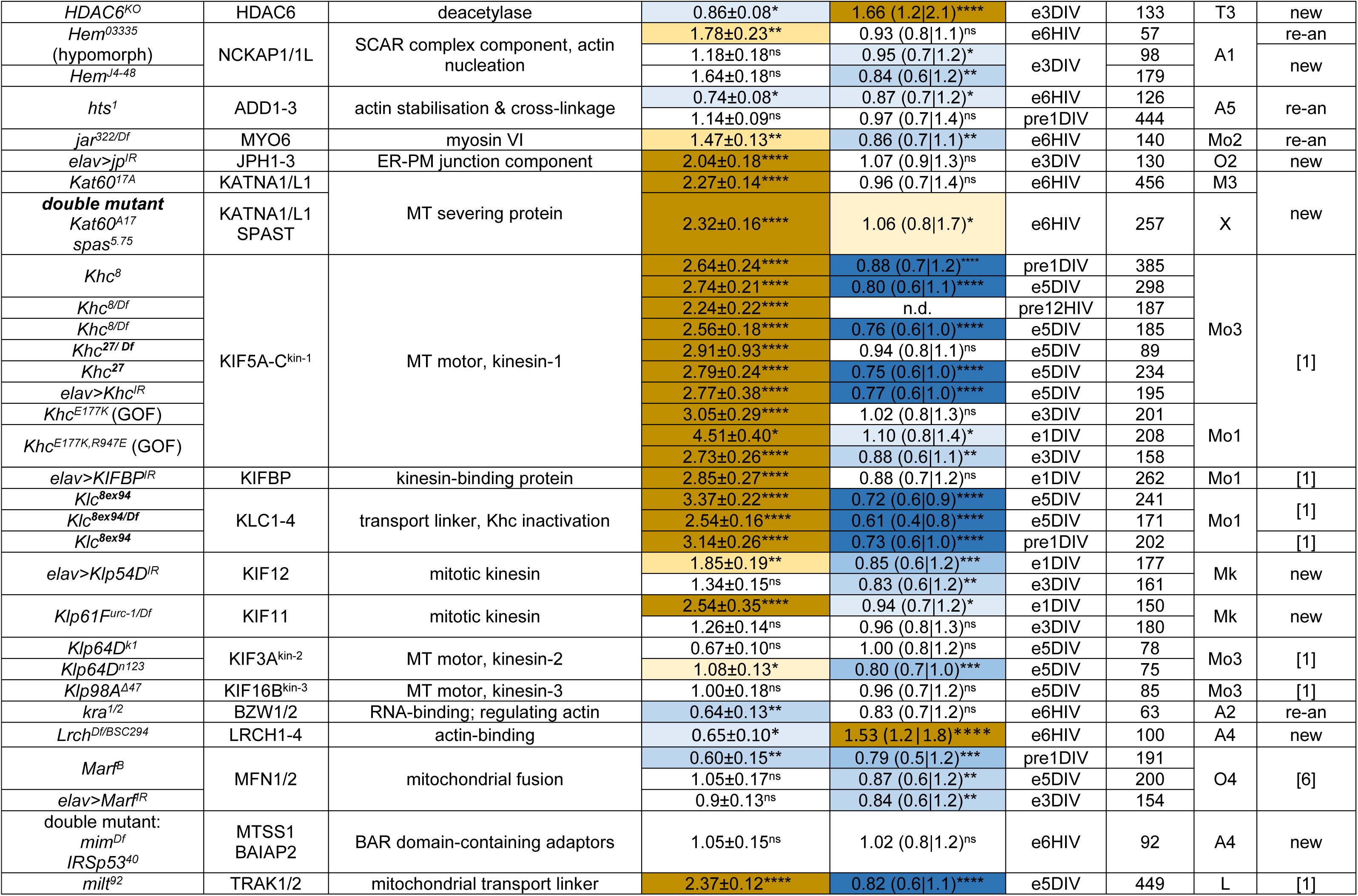
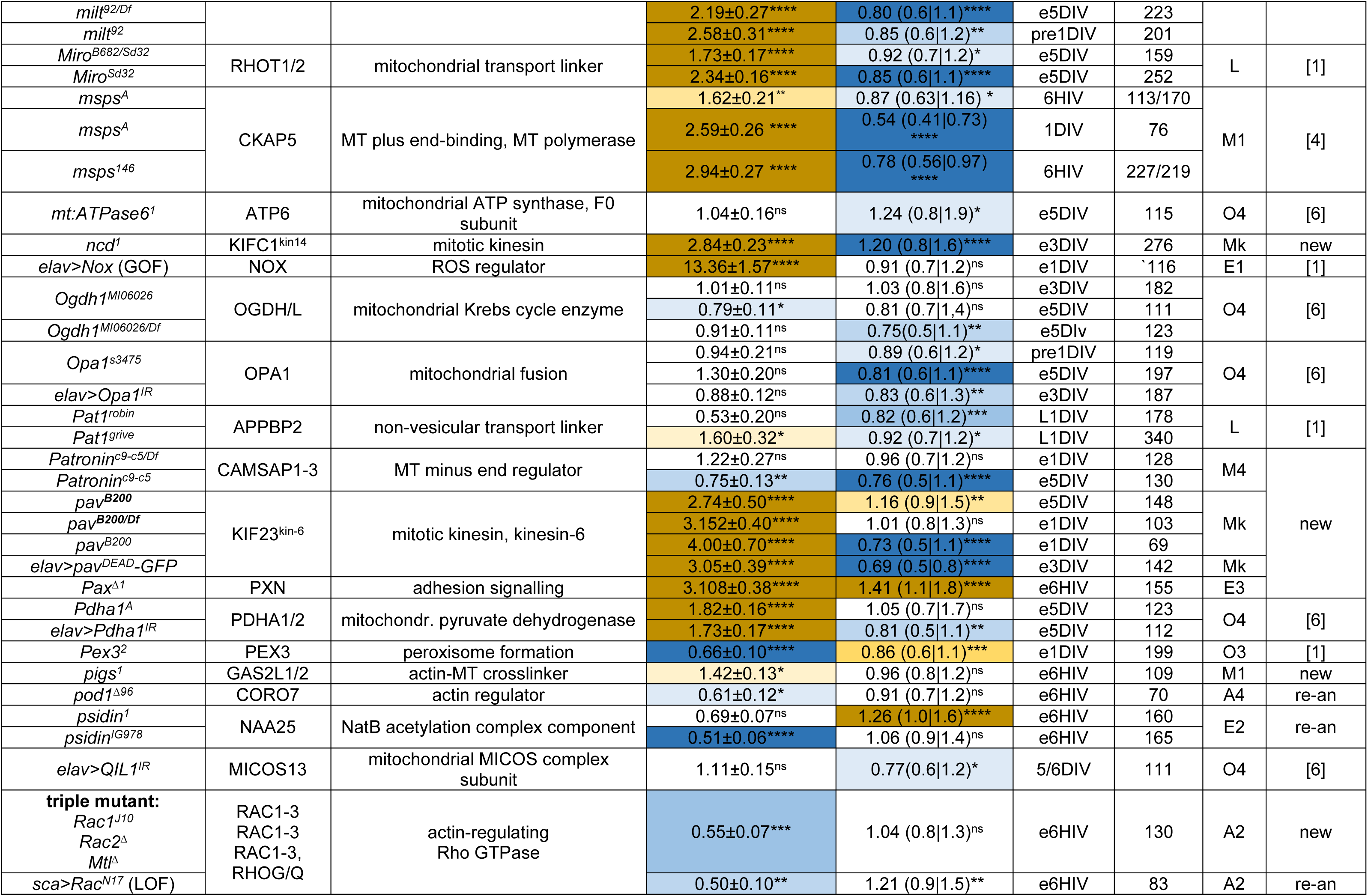
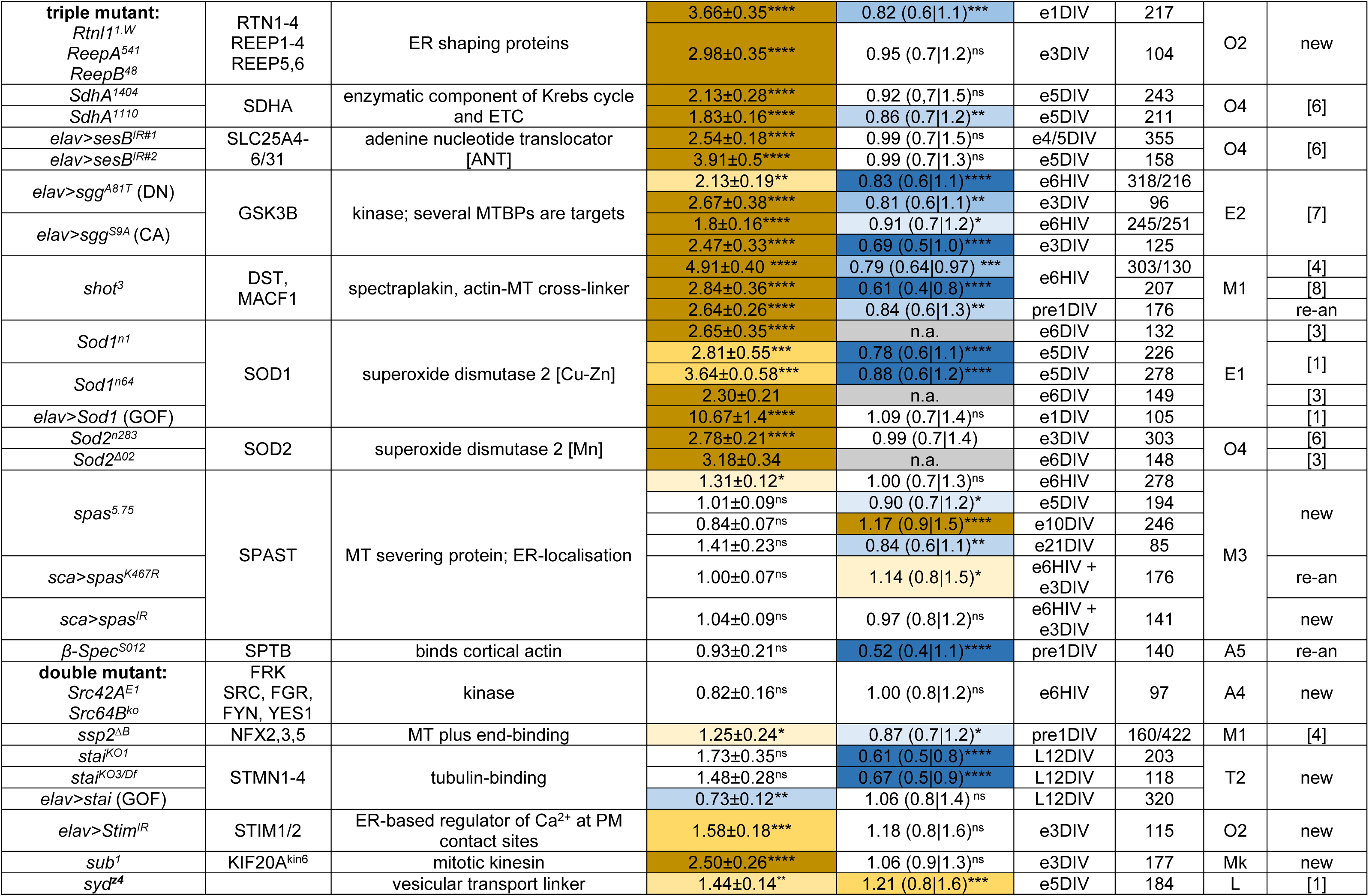
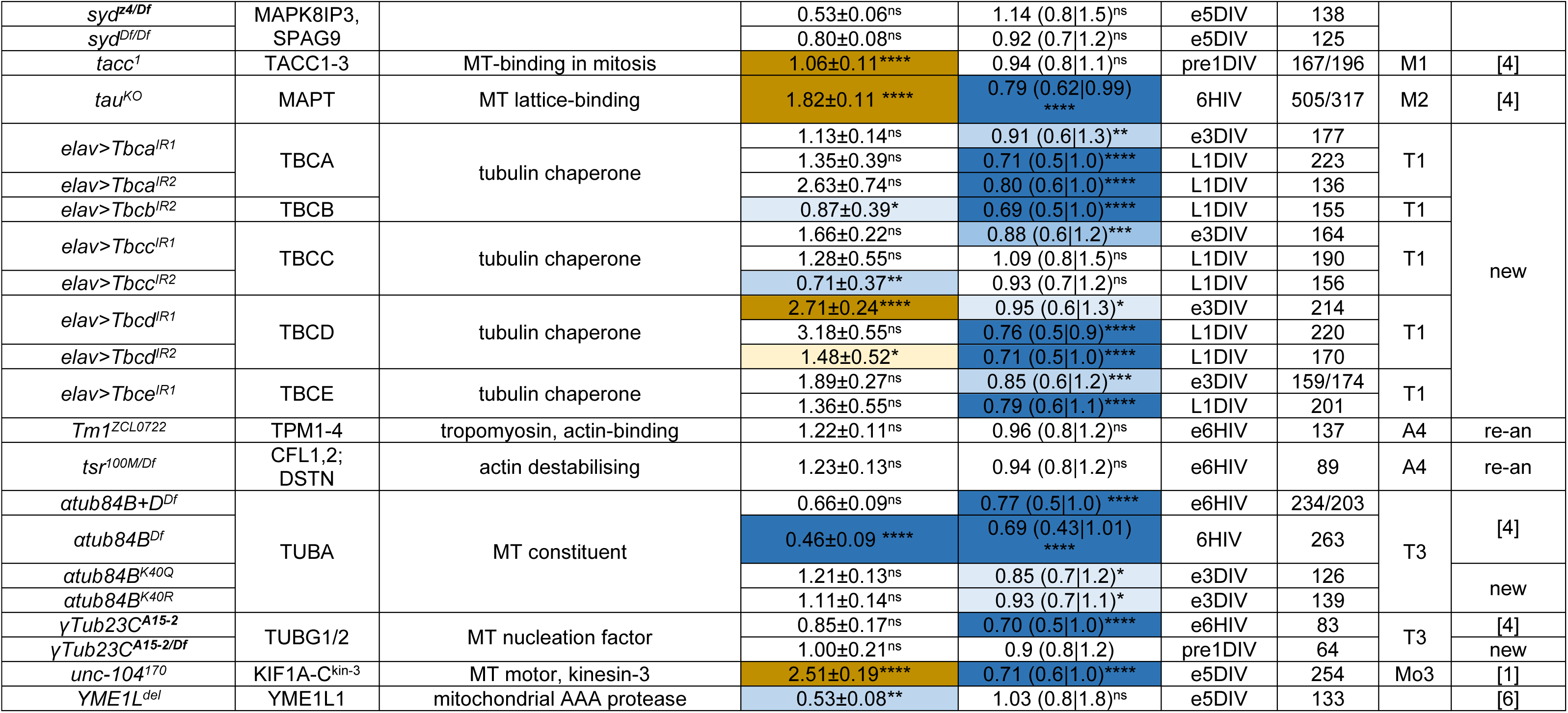
Genetic loss-of-function conditions assessed for axon length and MT-curling in *Drosophila* primary neuron cultures. **Column 1** shows the gene with its superscripted mutant allele name (alleles in hetero-allelic conditions separated by slash) or conditions where genes were knocked down (-IR) or overexpressed using Gal4 lines (named before ‘>’); GOF indicates gain-of-function conditions. **Column 2** lists corresponding close human orthologues as listed in flybase.org. **Column 3** provides a crude functional description of the respective gene. **Column 4** indicates the degree of curling given as mean ± standard error of mean for the assessed MT disorganisation index (MDI) normalised to internal wild-type controls; significance values established with Mann-Whitney U tests are indicated by colour (brown-orange/blue respresents increased/decreased curling); colour intensity matches the degree of significance indicated by superscript (ns, not significant P > 0.05; * P ≤ 0.05; ** P ≤ 0.01; *** P ≤ 0.001; **** P ≤ 0.0001; see Fig.1 for a colour code chart). **Column 5** shows the degree of axon length changes (median with quartile ranges) following the same colour and superscript concept as in column 4.**Column 6** informs about the respective culture condition: e, embryo-derived; DIV, days *in vitro*; HIV, hours *in vitro*; pre, embryo-derived and pre-cultured; L, derived from late larval CNSs. **Column 7** shows the number of neurons analysed (see also Methods). **Column 8** indicates which group (orange-highlighted code in Fig.1A) the gene was assigned to. **Column 9** indicates whether data are derived from unpublished experiments (‘new’), re-analyses of published experiments to a common standard (‘re-an’) or taken from the following previous publications: **[1]** (Liew et al., 2026); **[2]** (Gupta et al., 2026); **[3]** [Shields, 2025 #13494}; **[4]** (Hahn et al., 2021); **[5]** (Qu et al., 2019); **[6]** [Murray-Cors, 2025 #13899}; **[7]** (Voelzmann et al., 2026); **[8]** (Qu et al., 2022). Raw data files are available on figshare.com (https://doi.org/10.6084/m9.figshare.33112271).

**Mov.S1** Live-imaging examples of growing neurons and their MT-curling area dynamics. The movie shows two examples of wild-type, one *shot^3^* and one *Eb1^04524^*mutant neuron. Asterisks indicate cell bodies, arrow heads the growing axon tips, curved arrows areas of MT-curling; time of respective neurons is culture is indicated top left. Note that areas of MT-curling stay contained in wild-type but not in the mutant neurons. Movie available on figshare.com (https://doi.org/10.6084/m9.figshare.33112271).

